# Revealing ecologically coherent population structure of uncultivated bacterioplankton with POGENOM

**DOI:** 10.1101/2020.03.25.999755

**Authors:** C Sjöqvist, LF Delgado Zambrano, J Alneberg, AF Andersson

## Abstract

**Background:** Bacterioplankton are main drivers of biogeochemical cycles and important components of aquatic food webs. However, difficulties in culturing the majority of aquatic prokaryotic species have complicated the study of their microdiversity. Here, we present POGENOM, a software that quantifies population genomic indices from metagenome data, enabling comparative analysis of genomic diversity and differentiation in multiple species in parallel. We demonstrate POGENOM on metagenome-assembled genomes from the Baltic Sea and investigate their genomic variation using metagenome data spanning a 1700 km transect and covering seasonal variation at one station.

**Results:** The majority of the investigated species, representing several major bacterioplankton clades, displayed population structure correlating significantly with environmental factors such as salinity, temperature, nutrients and oxygen, both over horizontal and vertical dimensions. Population differentiation was more pronounced over spatial than temporal scales, although some species displayed population structure correlating with season. We discovered genes that have undergone adaptation to different salinity regimes, potentially responsible for the populations’ existence along the salinity range.

**Conclusions:** We provide a new tool for high-throughput population genomics analysis based on metagenomics data. From an evolutionary point of view, our findings emphasize the importance of physiological barriers, and highlight the role of adaptive divergence as a structuring mechanism of bacterioplankton species, despite their seemingly unlimited dispersal potential. This is of central importance when learning about how species have adapted to new environmental conditions and what their adaptive potential is in the face of Global Change.

## Background

Each litre of seawater contains around a billion bacterial and archaeal cells (bacterioplankton) that play central roles in biogeochemical cycles, marine food webs and ecosystem services ^1,2^. The diversity of aquatic prokaryotes is immense, by far exceeding that of eukaryotes. How this amazing diversity is generated and structured is not fully understood. Aquatic taxa are demonstrably differentially distributed among habitats ^3^ and ribosomal RNA (rRNA) gene sequencing has clearly shown that bacterioplankton communities are structured both in time and space and that the composition of operational taxonomic units (OTUs) is correlated with environmental parameters ^4–7^. While these studies have demonstrated that 16S rRNA gene clusters, or even specific 16S sequences (amplicon sequence variants; ASVs), represent organisms adapted to different habitats, the 16S rRNA gene does generally not provide enough genetic resolution to reveal within-species (intraspecific) diversity patterns, since prokaryotes with identical rRNA sequences may have highly divergent genomes and phenotypes ^8^. While comparative genomics of isolates, as well as metagenomics on natural samples, have revealed sequence clusters of >95% average nucleotide identity (which has emerged as an operational delineation of prokaryotic species ^9^), it is not known to what extent this intraspecific genomic variation represents neutral diversity vs. adaptation to different niches.

Due to the technical challenges, relatively little is known about intraspecific structuring of microbes, not least in the marine environment. However, pioneering studies have shown that genetic content of single bacterial species may correlate with geographic distance ^10^, and that coexisting but ecologically differentiated strains may arise through e.g. resource partitioning ^11^. Subtle differences in the phenotype, e.g. morphological and biochemical properties of cyanobacteria, have been linked to functional genes ^12^ and genetic clades or ‘ecotypes’ (<3% difference in 16S rRNA gene) of e.g. the cyanobacterium *Prochlorococcus* have been demonstrated to display niche differentiation along environmental gradients ^13^. The ubiquitous and most abundant type of organism in the ocean, the SAR11 clade, have undergone adaptive radiation in response to temperature ^14^. Apart from ‘genome streamlining’ ^14,15^, it is likely that the ecological success of this organism is facilitated by its adaptive divergence into ecotypes that are specialized for specific environmental conditions ^5,16^. A deeper understanding of intraspecific diversity, sometimes referred to as ‘microdiversity’ ^17^, is of crucial importance if we want to understand the ecology, evolution and speciation of bacterioplankton, and of prokaryotes in general. Studying genomic variation within a species can also reveal genes involved in adaptation to specific environmental factors, providing new clues on gene functions and cellular mechanisms of adaptation.

The study of metagenomes has been predicted to offer a more realistic view of prokaryotic diversity ^18,19^ as compared to PCR-based surveys of the rRNA gene. Metagenomics offers different routes for addressing intraspecific variation. The first is to reconstruct genetic information of individual strains. By mapping reads from one or several samples to the reference genome(s) of a species, the gene complement and/or nucleotide sequences at variant positions of the constituent strains can be inferred ^20–22^. This approach is promising, but challenging, especially in cases of many coexisting strains. The second approach does not aim at reconstructing strains, but rather uses the reads mapped to a reference genome to quantify intra- and intersample genomic variation of the species. This approach does not generate strain-resolved genomes but is more straightforward for analysing population structure and works well also in case of highly complex pan-genomes. Schloissnig et al.^23^ conducted pioneering cross-continental comparative analyses of human gut microbiomes using this approach and showed that there is more intraspecific genetic differentiation between habitats (human individuals) than within the same habitat over time. Similar approaches have been used by Nayfach et al. ^10^ and Delmont et al. ^24^ to show that gene content and amino acid composition, respectively, differ between oceanic regions within individual bacterial species.

Here we present the software POGENOM (POpulation GENOmics from Metagenomes) that, similar to MIDAS ^10^ and metaSNV ^25^, quantifies intraspecific genomic variation from metagenomic data. POGENOM differs from these software in that it takes as input a Variant Call Format (VCF) file, the standard file format for storing gene sequence variations. This allows the user to apply a variant caller of choice, rather than relying on an inbuilt algorithm. In addition, POGENOM automatically calculates a number of population genomic parameters for exploring both genome-wide and gene-specific patterns of variation.

We demonstrate the utility of POGENOM for the study of microdiversity within aquatic microbial species along natural gradients of salinity, temperature and nutrient concentrations in the Baltic Sea, covering a 1700 km S-N transect (Fig. 1a-c). This geologically young ecosystem is often used as a model for postglacial colonization, ecological differentiation and biodiversity clines ^26,27^. Marine macroorganisms display reduced species richness and intraspecific diversity towards higher latitudes in the Baltic region, as these impose more challenging conditions in terms of low salinity. Likewise, freshwater species diversity decreases with increasing salinity levels towards lower latitudes ^28^. Moreover, population genetic studies using have shown that species of fish and macroalgae have distinct genetic populations in the Baltic Proper (central Baltic Sea) compared to the Atlantic west of Sweden ^29–31^. With respect to unicellular organisms, population genetic data across the salinity regimes is only available for one eukaryote: the marine diatom *Skeletonema marinoi* ^32^. It is evident that the species is locally adapted and genetically differentiated into separate populations on each side of the Danish Straits, correlating with different salinity regimes and oceanographic connectivity. Herlemann et al. ^33^ studied bacterioplankton communities along the entire salinity gradient using the 16S rRNA gene and found the species composition to vary significantly along the salinity gradient, as well as vertically along oxygen gradients, with the Baltic Proper being composed of a mixture of typical freshwater and marine taxa. Using metagenomic binning and fragment recruitment analysis, Hugerth et al. ^34^ showed that the prokaryotic organisms in the Baltic Proper are genetically differentiated from closely related marine and freshwater lineages. However, it remains to be investigated whether the bacterioplankton populations are structured along the Baltic Sea environmental gradients. Here we use POGENOM to investigate population structure within multiple metagenome-assembled genomes (MAGs) across ten stations in the Baltic Sea, and also study population genomic patterns over time at one station.

**Figure 1.**
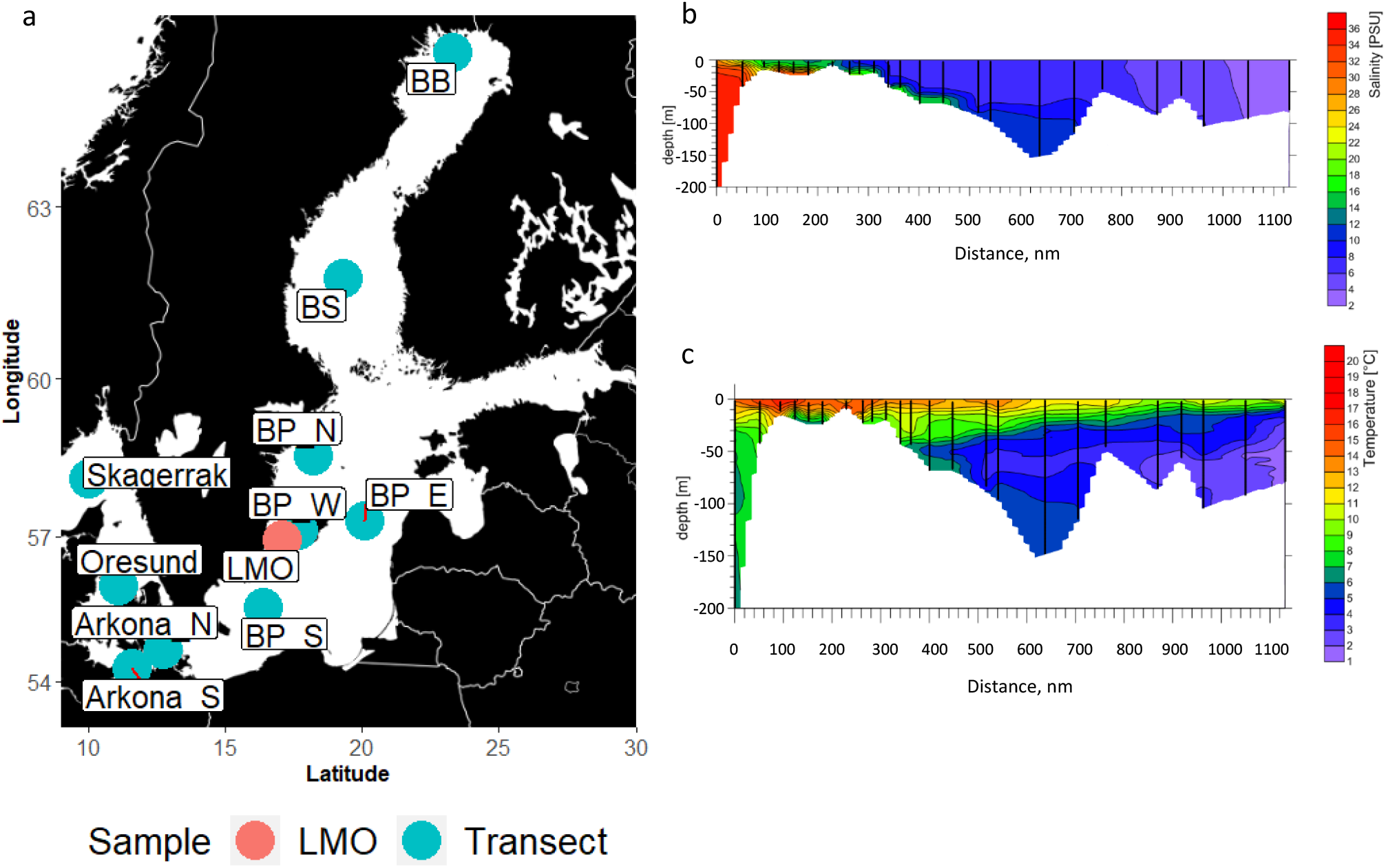
(**a**) The Baltic Sea. Sampling points marked with colored points. Transect samples (abbreviations in blue): BB=Bothnian Bay, BS=Bothnian Sea, BP_N=Northern Baltic Proper, BP_E=Eastern Baltic Proper, BP_W=Western Baltic Proper, BP_S=Southern Baltic Proper, Arkona_N=Northern Arkona basin, Arkona_S=Southern Arkona basin. At each point samples were retrieved from three different depth layers ^35^. Time series samples (abbreviation in red): LMO=Linnaeus Microbial Observatory. In total 31 samples were retrieved between March-December ^34^. (**b**) Water temperature across the horizontal transect with depth variation. The x-axis shows the distance from Skagerrak sample in nautical miles. (**c**). Salinity across the horizontal transect with depth variation. x-axis as in b.

We hypothesized that 1) the intra-sample diversity (π) of typical marine prokaryotic taxa (SAR11, OM182) should decrease towards low salinity and display the opposite trend for freshwater lineages (e.g. Actinobacteria), 2) population genomic structure quantified by the fixation index (*F*_ST_) should correlate with environmental gradients of salinity, temperature and various nutrient concentrations and show greater differentiation across space than time, and 3) individual genes should be under natural selection as a result of differing environmental regimes. With this approach, we aim at explaining patterns of intra-specific variation in aquatic prokaryotes and elucidate the selection pressures different subpopulations experience across the ecosystem.

## Results

In order to better understand patterns of intraspecific variation and population genomic structure in bacterioplankton, we applied our newly developed software POGENOM on a set of MAGs that was recently reconstructed from a time-series dataset at a station within the central Baltic Sea (the Linnaeus Microbial Observatory [LMO]) ^34^. The LMO MAG set includes 83 MAGs that were earlier clustered based on sequence identity into 30 species-level clusters (BAltic Sea CLusters; BACL) (Hugerth et al. 2015). For each BACL, one representative MAG was used in the analysis here (Table 1, Supplementary Table 1). Metagenome reads from 30 samples of the original LMO dataset were mapped to the 30 MAGs (BACLs). Reads were also mapped from 30 metagenomic samples obtained in a transect cruise ^36^, where 10 stations distributed along the Baltic Sea salinity gradient where sampled at three depths (Figure 1, Supplementary Table 2). To lower the risk of including reads derived from other species, we only included reads mapping with >95% identity to the MAGs. We set a median coverage depth threshold of ≥20X (Supplementary Table 3) and a minimum coverage breadth of 40% (Supplementary Table 4) to include a sample for a BACL. For each transect depth layer, and for the LMO time-series, only BACLs with ≥5 samples fulfilling these criteria were included in the downstream analyses. Using these criteria, we obtained ten BACLs from the transect data set and seven BACLs from the time-series data set eligible for further analysis.

**Table 1.** MAGs included in the population genomic analysis and their overall SNP frequency and mean within-sample nucleotide diversity (π).

| Baltic Cluster | MAG ID | SNP frequency | Intra-pi | Taxonomy |
| --- | --- | --- | --- | --- |
| BACL1 | NA | 0.0311 | 0.0029 | BACTERIA;PROTEOBACTERIA;GAMMAPROTEOBACTERIA;SAR86; |
| BACL2 | NA | 0.0139 | 0.0036 | BACTERIA;ACTINOBACTERIA;ACI; |
| BACL3 | NA | 0.0078 | 0.0016 | BACTERIA;PROTEOBACTERIA;GAMMAPROTEOBACTERIA;OM182; |
| BACL5 | NA | 0.0035 | 0.0010 | BACTERIA;PROTEOBACTERIA;ALPHAPROTEOBACTERIA;SAR11; |
| BACL6 | NA | 0.0089 | 0.0021 | BACTERIA;ACTINOBACTERIA;ACIV; |
| BACL8 | NA | 0.0025 | 0.0005 | BACTERIA;BACTEROIDETES;FLAVOBACTERIIA;FLAVOBACTERIACEAE; |
| BACL10 | NA | 0.0113 | 0.0021 | BACTERIA;PROTEOBACTERIA;ALPHAPROTEOBACTERIA;RHODOBACTER; |
| BACL11 | NA | 0.0113 | 0.0030 | BACTERIA;BACTEROIDETES;FLAVOBACTERIIA;CRYOMORPHACEAE; |
| BACL14 | NA | 0.0033 | 0.0011 | BACTERIA;PROTEOBACTERIA;BETAPROTEOBACTERIA;OM43; |
| BACL15 | NA | 0.0033 | 0.0009 | BACTERIA;ACTINOBACTERIA;ACI; |
| BACL20 | NA | 0.0092 | 0.0022 | BACTERIA;PROTEOBACTERIA;ALPHAPROTEOBACTERIA;SAR11; |
| BACL27 | NA | 0.0036 | 0.0012 | BACTERIA;ACTINOBACTERIA;ACIV; |
| BACL28 | NA | 0.0087 | 0.0026 | BACTERIA;ACTINOBACTERIA;LUNA; |
| BACL30 | NA | 0.0118 | 0.0031 | BACTERIA;CYANOBACTERIA;CYANOBIUM; |

### Nucleotide diversity

In total, 171,527 single-nucleotide polymorphisms were identified in the 14 BACL genomes, with frequencies ranging from 2.5 kbp^−1^ (BACL8) to 31 kb^−1^ (BACL1). Mean within-sample nucleotide diversity (π), corresponding to the likelihood that two metagenome reads that overlap a position in the genome will differ at the position, ranged from 0.00054 (BACL8) to 0.0036 (BACL2) (Table 1). We only found a few significant correlations between π of different BACLs and environmental variables subsequent to post-hoc tests across the Baltic Sea transect (Fig. 2a, Supplementary Table 5), contradicting our first hypothesis. For instance, total nitrogen explained the variation in nucleotide diversity for BACL6 across the Baltic Sea, but we observed no other correlations between π and environmental variables in the surface layer. The level of diversity over time was also remarkably stable (Fig. 2c). However, the genome-wide nucleotide diversity in BACL1 (belonging to the SAR86 clade) correlated with season and temperature. This supports previous findings ^7^ showing a temperature-dependent distribution of this uncultivated Gammaproteobacterial clade. When the surface layer and time series data were included in the same analysis, significant correlations appeared with salinity, chlorophyll *a* and DOC.

**Figure 2.**
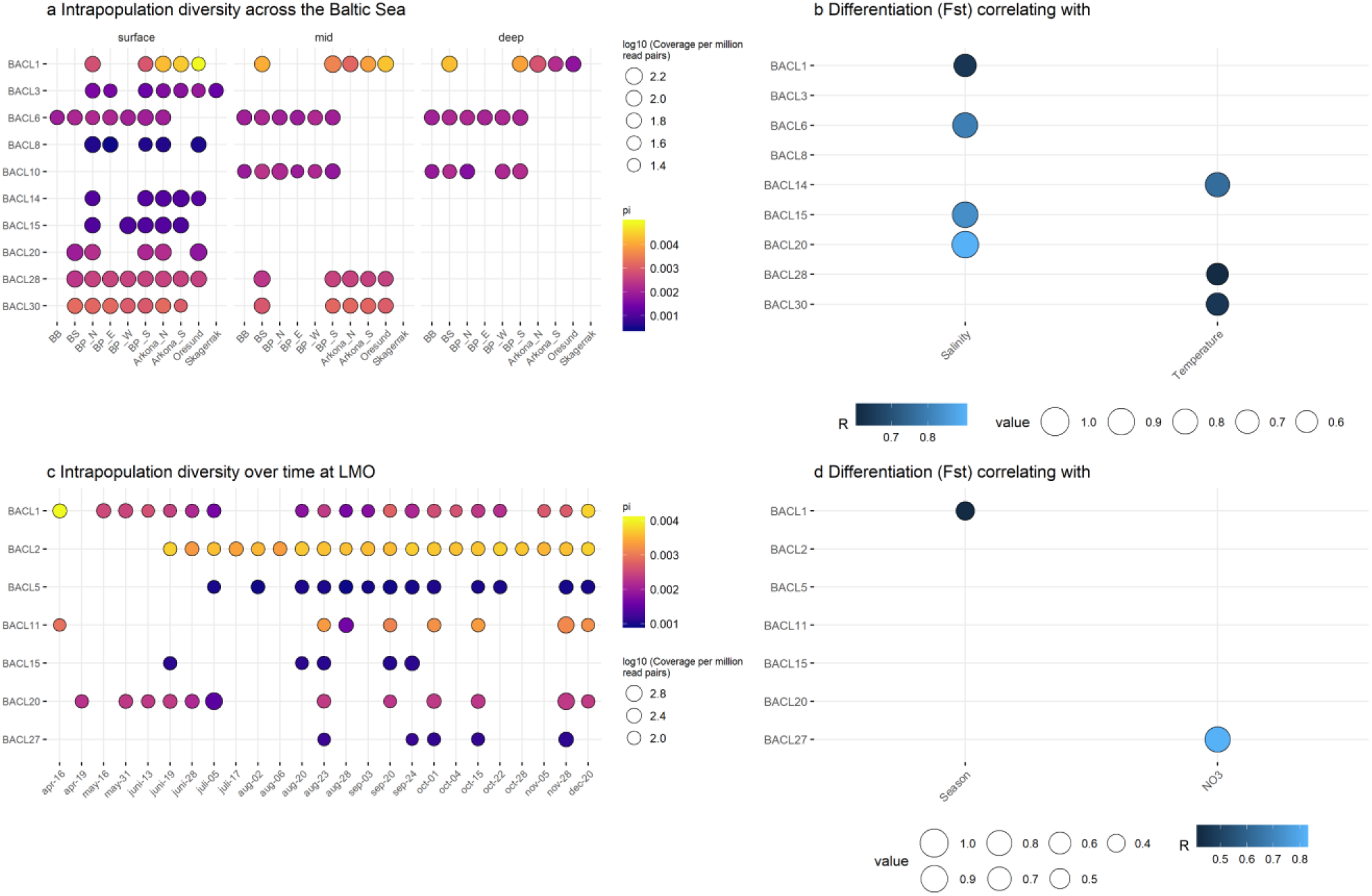
(**a**) Nucleotide diversity (π) for BACLs across the Baltic Sea transect. (**b**) Environmental variables (single best predictor) correlating with population genomic differentiation among the transect surface samples. (**c**) Nucleotide diversity (π) for BACLs over time at the LMO station (note different scale in a and c). (**d**) Environmental variables (single best predictor) correlating with population genomic differentiation among the LMO samples. Only significant correlations are shown in b and d (FDR adjusted *P*-value < 0.05).

The finding that intra-population diversity was largely unchanged along the salinity gradient contrasts to the situation in macroorganisms, where many species have been reported to display variation in diversity between different basins of the Baltic Sea ^27^.

POGENOM also outputs nucleotide diversity per gene. As expected, this was in most (10/14) BACLs on average smaller for housekeeping genes compared to other genes ^23^, using a set of 36 single-copy core genes (SCGs) that are found once in almost all known bacterial genomes ^37^. The difference reflects that the housekeeping genes are under stronger purifying selection than an average gene, and was even more pronounced when instead comparing non-synonymous to synonymous polymorphism rates (pN/pS) ^38^ (Supplementary Figure 1).

POGENOM outputs per-sample allele frequencies both at the nucleotide level (SNV) and, for protein-coding regions, at the amino acid level (single amino acid variations; SAVs). Halophilic prokaryotes that employ the “salt-in” strategy, i.e maintain high intracellular salt concentrations, have fundamentally adapted their proteome to be compatible with this. These include the presence of a large excess of acidic amino acids and small amounts of hydrophobic amino acids ^39^. Of the nine BACL analysed for the transect surface water, only one (BACL1) displayed a clear pattern of increasing frequencies of acidic amino acids and decreasing frequencies of alkalic amino acids with increasing sample salinity (Supplementary Figure 2).

### Population genomic structure

Of the nine BACLs that were present in at least five of the surface transect samples, five displayed a significant correlation between fixation index (*F*_ST_) - a measure of population differentiation across samples - and salinity level (Spearman False Discovery Rate-adjusted *P*-value (*Q*) < 0.05) (Fig. 2b, Supplementary Table 5) when only looking at the single best predictor of population genomic structure. Temperature was another significant driver of population structure in the surface water (BACL14, BACL28). These results show that aquatic bacterial species may diverge from the null hypothesis of panmixia and that populations are structured by species-specific environmental drivers. This implies the existence of ecotypes ^40^ that may remain undetected by rRNA gene sequencing. Including combinations of independent variables in the analysis revealed that the population genomic structure of Baltic Sea bacteria is mainly driven by a combination of salinity and temperature (Supplementary Table 6). In some cases there appears to be isolation by distance, as the MEM (Moran’s eigenvector maps) variables (spatial factor) show significant correlation with genome-wide *F*_ST_ in the conditioned RDA analysis (i.e. controlling for the explanatory power of all other included variables, see *Environmental Association Analysis* in Methods).

Of the seven BACLs that were present in at least five of the LMO samples, only one (BACL1) displayed a significant correlation between *F*_ST_ and season (Spearman *⍴* = 0.42, *Q* < 0.05) (Supplementary Table 6). When more than one explanatory variable was allowed, a combination of temperature and season explained the variation in *F*_ST_ over time for BACL1 (global RDA, *R^2^* adj. = 0.61, *P* = 0.001). In the conditioned RDA (controlled for ‘season’), temperature showed a stronger individual effect (*R^2^* adj. = 0.40, *P* = 0.001) than season (*R^2^* adj. = 0.20, p = 0.002). The population genomic structure of BACL27 appears to be driven by variation in nitrate levels at the LMO station (*⍴* = 0.82, *Q* < 0.05). *F*_ST_ for BACL15 correlated significantly with chlorophyll *a* as shown in the RDA-analysis (Supplementary Table 6), implying an association of the species to the succession of the phytoplankton community.

In the BACLs for which a comparison between genomic differentiation over time and space was feasible, we found that the magnitude of differentiation was greater spatially across the Baltic Sea than over time at the LMO station (Table 2). Hence, our data show that genomic differentiation is more prevalent over the spatial than the temporal scale in the Baltic Sea ecosystem, analogous to what has previously been proposed for species sorting at the community level ^41^. Thus, the diverse and environmentally structured bacterioplankton communities that have been observed earlier ^33^, seem to be even further differentiated at the species level.

**Table 2.** Comparison of *F*_ST_-values across the Baltic Sea vs. over time at station LMO. Mean and max *F*_ST_ values are reported as well as *P*-values from Wilcoxon rank-sum tests comparing the distributions of *F*_ST_ values over time (LMO) and space (Transect).

| MAG | mean Fst (LMO) | max Fst (LMO) | mean Fst (Transect) | max Fst (Transect) | Wilcoxon p-value | n |
| --- | --- | --- | --- | --- | --- | --- |
| BACL1 | 0.144 | 0.479 | 0.365 | 0.737 | $>0.0001$ | 295 |
| BACL15 | 0.114 | 0.142 | 0.172 | 0.265 | 0.001 | 20 |
| BACL20 | 0.156 | 0.676 | 0.271 | 0.505 | 0.002 | 88 |

**Table 3.** Correlations between gene-wise *F*_ST_ and difference in salinity among the surface layer transect samples. “#Genes correlated” gives the number of genes displaying a Spearman correlation with FDR-adjusted *P* < 0.05. “#Genes with higher correlation than expected” gives the number of genes with higher correlation than expected (FDR-adjusted *P* < 0.1), given the genome’s background correlation (based on 1000 permutations). “#Included” gives the number of genes included in the analysis.

| MAG | #Genes correlated | #Genes with higher correlation than expected |  |
| --- | --- | --- | --- |
| | ($p < 0.05$) | ($q < 0.10$) | (#included) |
| BACL1 | 92 | 14 | 1293 |
| BACL3 | 489 | 46 | 1371 |
| BACL6 | 406 | 28 | 1266 |
| BACL8 | 10 | 0 | 516 |
| BACL14 | 1 | 0 | 534 |
| BACL15 | 5 | 5 | 498 |
| BACL20 | 49 | 0 | 410 |
| BACL28 | 14 | 17 | 763 |
| BACL30 | 42 | 56 | 1613 |

When samples from both the transect and LMO were included in the environmental association analysis, salinity and temperature were the most important drivers of population genomic structure. *F*_ST_ of five BACLs were also compared across the vertical dimension, i.e. water depth. In three cases (BACL1, BACL28, BACL30), oxygen was the single best predictor of population genomic structure across depth. However, a combination of oxygen, salinity and geographic location provided even better explanatory power for variations in *F*_ST_ for BACL1. In BACL10, a combination of temperature and geographic location was statistically significant, however, variation partitioning showed that temperature was more important in driving the observed differences in this BACL.

Two BACLs were represented in all three depth layers in the transect sample set (BACL1 and BACL6). Principal Coordinate Analysis (PCoA) based on the *F*_ST_ values visualizes the population structure of these BACLs over horizontal and vertical scales (Fig. 3a-b). For both BACLs, salinity correlated significantly with the first principal coordinate and depth with the second (Spearman *P* < 0.01; Fig. 3b,c,e and f). According to the global RDA analysis using both horizontal and vertical data, salinity and oxygen correlated significantly with genomic differentiation for BACL1 (*R^2^*-adj. = 0.75, *Q* < 0.001) (Supplementary Table 6). This demonstrates that population structure of bacterial species can be driven by different environmental variables across horizontal and vertical scales. BACL1 belongs to the SAR86 clade, a ubiquitous Gammaproteobacteria occurring in high abundances in oceanic surface waters ^42,43^. Several phylogenetic subgroups within the SAR86 clade have been observed, but it has remained largely unknown if the differentiation is related to variation in environmental factors ^44^.

**Figure 3.**
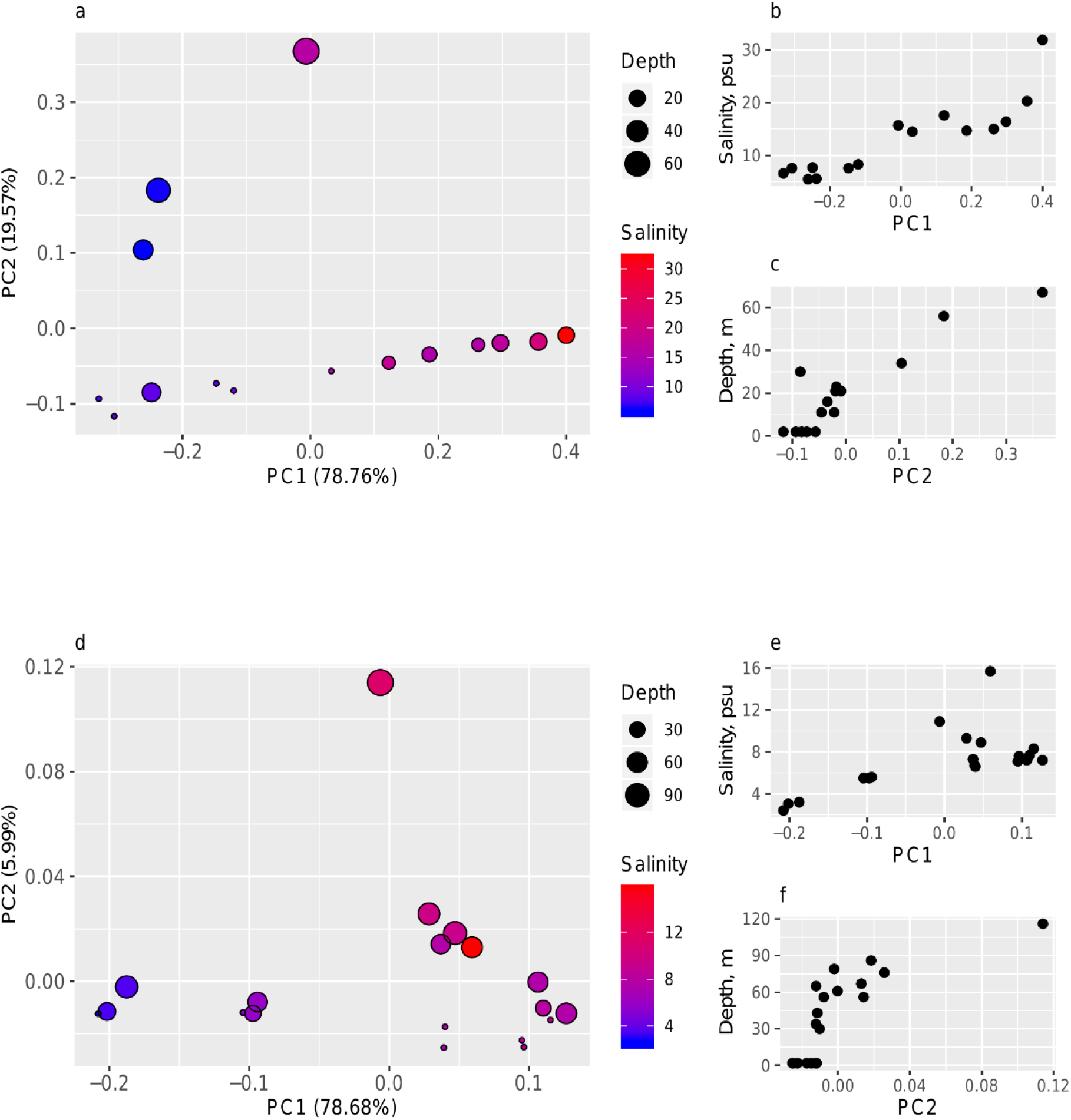
(**a**) PCoA of the population structure in BACL1 and BACL6 based on *F*_ST_ values. Samples are represented by circles. Color and size of the circles correspond to salinity and sampling depth, respectively. Samples proximal in the graph display low genomic differentiation between corresponding populations. (**b**) Scatterplot of Principal Coordinate 1 (PC1) in BACL1 vs. salinity. (**c**) Scatterplot of PC2 in BACL1 vs. water depth. (**d,e**) Same as (**b,c**) but for BACL6.

### Population structure at the gene-level

The results above show that most populations are structured according to environmental factors when fixation index is measured at the whole-genome level. However, such results may emanate from distinctive differentiation from merely a few genes or genomic segments. To facilitate the interpretation of population structure, POGENOM also allows analysing *F*_ST_ at the individual gene level. This revealed that a number of genes had *F*_ST_ values that were positively correlating with difference in salinity for the transect surface samples (Table 2). These correlations could indicate that the genes themselves have undergone adaptation to the different salinity levels or to environmental factors that co-vary with salinity. Alternatively, it could reflect genetic hitchhiking, i.e. that an allele that differs in frequency between environmental conditions do so, not because it itself has been under natural selection, but because other gene(s) on the same genome have undergone selective sweeps: if the rate of homologous recombination is low relative to selection, unrelated alleles in the genomes of strains carrying the beneficial gene (or gene variant) will also increase in abundance in the population ^45^. To differentiate between these processes, we devised a permutation procedure, where permuted *F*_ST_ values are calculated for each gene, while keeping the population differentiation constant at the genome-level. Basically, all variant loci are shuffled for the genome so that each gene will get a new set of loci (with their associated allele frequencies) while having the same number of variant loci as in the original data. Subsequently, gene-wise *F*_ST_ values are calculated for all pairs of samples. The process is repeated many times and the permuted *F*_ST_ values can then be compared with the original. This procedure is advantageous to other modelling approaches in that no assumptions on population histories are needed.

The permutation procedure indicated that a fraction of the genes indeed had *F*_ST_ values that correlated stronger with salinity than expected from the genome background level (Table 2). Such genes were evident in BACLs with population structure correlating with salinity at the whole genome-level (such as BACL3 and 6), but interestingly also in two of the BACLs that did not display such correlations (BACL28 and 30). Looking at the distribution of correlations between gene-wise *F*_ST_ and salinity difference revealed a small number of genes in these genomes with markedly higher correlation than expected by chance (Fig. 4), indicating that these genes have undergone adaptive evolution and may be partly responsible for that the populations can exist along a range of salinity levels. It is possible that high homologous recombination rates in these populations result in selective sweeps acting more at the gene than at the genome level.

## Discussion

We estimated genome-wide diversity and population differentiation for a set of uncultured aquatic prokaryotic species along environmental gradients across the Baltic Sea. We quantified population genomic indices, such as the intra-population diversity (π) and the fixation index (*F*_ST_), and also studied potential selection on a per-gene basis by computing gene-wise *F*_ST_ and comparing with gene-wise *F*_ST_ obtained after randomly redistributing variant loci across the genome. With these analyses, we obtained information about environmental drivers of population structure and indications on individual genes under selection to different environmental regimes. Such an exercise is now significantly streamlined with the advent of POGENOM, calculating all the above parameters automatically. With intra-population diversity (π) we refer to the average nucleotide diversity of a population. The fixation index (*F*_ST_) measures the differences in allele frequencies between pairwise populations and takes a value between 0-1. *F*_ST_ was originally designed for diploid, sexually reproducing organisms ^46^ where a value close to 1 is interpreted as substantially restricted gene flow. However, the concept of *F*_ST_ involves no obligate condition of sexual reproduction, as it simply compares allele frequencies between two populations and is thus as valid for asexually reproducing prokaryotes ^47^. The reasons for an observed value of *F*_ST_ between pairwise prokaryotic populations only needs to be interpreted with the biological facts in mind. For instance, constraints in gene flow as apprehended in sexually reproducing organisms may for prokaryotes be seen as constraints in homologous recombination and/or effects of environmental sorting of genetic material, leading to skewed allele frequencies between populations. Shapiro ^48^ showed that ecological differentiation of bacterial species may, in fact, be akin to the process in sexually reproducing eukaryotes. The main process involves adaptive mutations sweeping through subpopulations, initiating specialisation to different habitats and thereby limiting gene flow between subpopulations, leading to gradually increasing genetic separation. It is, however, worth keeping in mind that pure chance may play an important role in the genomic makeup of populations. Especially when these are physically separated and have small effective population sizes, genetic drift, and not natural selection, may be the major cause of differences in allele frequencies ^49^. However, bacterioplankton have large effective population sizes and readily migrate, suggesting that the most likely driver of genetic differentiation in distinct loci is natural selection (Kashtan et al. 2014).

Our analyses showed that the majority of the BACL had a genomic population structure significantly correlating with environmental variables. For surface waters, in particular with salinity and temperature. Several BACLs also displayed statistically significant isolation by distance (IBD; Supplementary Figure 3), similar to the findings in Nayfach ^10^, but variation partitioning analyses indicated that the geographic effects were generally not significant after taking environmental factors into consideration (Supplementary Table 6). In cases where stable differential selection is sustained, as in the Baltic Sea, and where geographic distances are rather small, isolation by adaptation (IBA) can spur such population genomic structure within a species ^50,51^. Our results suggest that bacterioplankton do not only modulate the flexible gene pool when confronted by environmental clines, as demonstrated by several studies ^10,52,53^, but also adapt by modifying existing genes (altering allele frequencies) to the different environmental conditions. Delmont ^24^ suggested an evolutionary mechanism for such conservation of genetic heterogeneity by emphasizing the role of consistent purifying selection against deleterious non-synonymous variants, exemplified by the cosmopolitan SAR11 clade. The same authors showed a partitioning of SAR11 metagenomes in concordance with large-scale oceanic temperatures suggesting that environmental selection is of central importance even at the microdiversity level in marine bacterioplankton. We demonstrate a similar example of environmentally driven population structure in e.g. one Actinobacteria species (BACL6) belonging to the acIV clade along the transect from the Bothnian Bay to the southern Baltic Proper, suggesting its population genomic structure was mainly driven by salinity in this system (*ρ* = 0.80, *Q* < 0.05) (Supplementary Table 6). Actinobacteria are one of the most abundant types of freshwater bacteria and comprise multiple different clades and species-level clusters ^54,55^. Recent discoveries on their marked microdiversification may explain their success in environmentally variable conditions ^56^. The Baltic Sea also hosts an abundant community of different cyanobacteria species. An example is the cosmopolitan genus *Synechococcus*, which is prevalent in the Baltic Sea during summer or when the temperature reaches >15°C ^57^. The environmental association analysis showed that the population structure of the *Synechococcus sp*. analysed here (BACL30) was mainly driven by temperature in the surface layer (*ρ* = 0.64, *Q* < 0.01). This BACL was also differentiated across depth, which is in line with earlier findings of different pigmentation phenotypes utilizing a variety of light wavelengths ^58^. When analyzed across depth layers, we also found a significant correlation between population structure and oxygen (*ρ* = 0.37, *Q* < 0.05). Szabo ^59^ found high reproducibility of coastal bacterioplankton population genetic structure by observing the *hsp*-gene in four *Vibrionaceae* clusters over time, implying the population structure is non-random. Our analyses show that the population genomic structure over time correlated significantly with fluctuations in temperature (BACL1) and NO_3_-levels (BACL27) (Supplementary Table 6), suggesting high micro-niche fidelity in some of the analyzed genomes. The theory of ecotypes, defined as “populations that are genetically cohesive and ecologically distinct” ^40^ builds on the same type of dynamics. Earlier comparative genomic studies have shown remarkable differences in gene content of bacterioplankton belonging to the same species and concluded that the flexible part of the genome is modified by horizontal gene transfer as a response to selective forces ^60,61^. For example, Nayfach^10^ found population structure, in terms of gene content, in marine bacteria to correlate with geography, but to our knowledge, this is the first study to show explicitly how allele frequencies in marine bacterial populations are dictated by species-specific environmental drivers.

The lack of pattern in intra-sample diversity (π) along the salinity gradient contrasts to the situation reported in macroorganisms, where, at least for species of fish and macroalgae, π varies along the salinity gradient ^27,62^. However, our previous study based on MAGs from station LMO and fragment recruitment of globally distributed aquatic metagenomes indicated that the bacterioplankton of the Baltic Sea are members of a globally distributed brackish metacommunity, rather than locally adapted freshwater and marine taxa. Thus, unlike most macroorganisms in this ecosystem, the planktonic prokaryotes residing in the Baltic Sea were likely adapted to brackish conditions already when they entered the system. Whether most of the intraspecific variation and niche differentiation that we see within the different BACLs was gained after the populations immigrated, or were in place already before, as a set of strains with different genetic make-up and ecological niches, we cannot give a conclusive answer to here. However, it is interesting to note that some populations display gene-wise *F*_ST_ correlations to environmental gradients for a large number of genes, while others only do it for a few. This may reflect that the populations of the latter have adapted more recently to the different environmental conditions (possibly after colonising the Baltic Sea) via selection only on the most critical genes. Recently, proteome differences between some freshwater prokaryotes and their closest marine relatives could be observed, with a larger proportion of acidic and a lower number of alkalic amino acids in the proteome of the marine representative of each pair, indicating that adaptations changing the chemical properties of the proteome may be important for crossing the freshwater - marine boundary ^63^. We only observed such a pattern in BACL1, suggesting that adaptations altering the physicochemical properties of the proteome are not the major driver behind the population structure that we observe within the Baltic Sea region for these populations. However, it may have been important for facilitating the transition from freshwater or marine to brackish conditions in the first place.

## Conclusions

Facilitated by our recently developed program POGENOM, we show that populations of multiple bacterioplankton clades are genomically structured, even within the same ecosystem. Genomic differentiation of populations correlated with environmental variables such as salinity, temperature and nutrient levels across spatial dimensions and in some cases also over time. This emphasises the role of isolation by adaptation rather than isolation by distance as a driving force for speciation of aquatic prokaryotes. Population genomics analysis based on metagenomics data will undoubtedly lead to a deeper understanding of the ecology and evolution of important and uncultivated bacterioplankton species. We show that POGENOM may advance the understanding of microdiversity in bacterioplankton which is of central importance when learning about how species have adapted to new environmental conditions and what their adaptive potential is in the face of Global Change.

## Methods

### POGENOM software

POGENOM takes as minimal input a file of the variant call format (VCF). This is generated by mapping one or several metagenome samples against a reference genome with a read aligner such as Bowtie2 ^65^, BWA ^66^ or MOSAIK ^67^ and calling variants using a variant caller such as GATK ^68^ or Freebayes ^69^. POGENOM calculates the nucleotide diversity (π) within each sample. If multiple samples have been mapped, the fixation index (*F*_ST_) is calculated for all pairs of samples. If, in addition to the VCF file, an annotation file of the General Feature Format (GFF) is provided, gene-wise π and *F*_ST_ will be calculated. If further, the genome sequence is provided in the GFF file or in a separate FASTA file, amino acid frequencies will be calculated for each codon position in each gene and sample, and gene-wise π and *F*_ST_ will be calculated also at the amino acid level. Now also non-synonymous to synonymous polymorphism rates (pN/pS) will be calculated for each gene and sample. POGENOM has several optional parameters, such as minimum read depth for a locus to be included for a sample, minimum number of samples with minimum read depth for a locus to be included at all, subsampling to a given read depth, splitting of haplotypes into individual SNVs in case haplotype variant calling was applies, etc. A complete description on how the different parameters are calculated can be found in the Supplementary Information. POGENOM is implemented in Perl, its source code and documentation and a pipeline for automatic generation of input data (VCF files) are available at https://github.com/EnvGen/POGENOM.

### Sampling, library preparation and sequencing

Metagenomes from the 30 transect samples were included from a previous study ^35^. Briefly, water from three different depth layers across the Baltic Sea was retrieved using a compact CTD profiling instrument (HYDRO-BIOS, Kiel, Germany) and samples for DNA analysis were captured on 0.2 μm pore size filters (GVWP04700, Merck Millipore, Darmstadt, Germany) and DNA extracted and stored in −80°C until further processing. Shotgun library preparation and sequencing were conducted at the National Genomics Infrastructure (NGI) at Science for Life Laboratory, Stockholm, Sweden, using a full HiSeq 2500 high-output flowcell, which generated on average 69.5 million paired-end reads per sample. We also included time-series samples from a previous study ^34^ where water samples were collected with a Ruttner sampler at the LMO station (Linnaeus Microbial Observatory; 56°55.851, E 17°03.640) in the central Baltic Sea. This station was sampled 1-6 times per month from March 2012 to December 2012 resulting in 31 water samples (Supplementary Table 2). The time-series samples were filtered onto 0.2 μm pore size filters after pre-filtration through 3.0 μm pore size filters. Shotgun sequencing on the extracted DNA was conducted as described above. This set of samples generated on average 31.9 million paired-end reads per sample.

### Mapping of sequencing data

In total 66 (30 transect + 31 time series) quality filtered samples (as in ^34^) were mapped against 30 previously generated metagenome-assembled genomes (MAGs) ^34^ using MOSAIK ^67^. Each MAG belongs to a different BAltic Sea CLuster (BACL) and can be considered different species ^34^. The similarity of MAGs within a BACL is >99%. We mapped against the genome with the highest N50 length value within each BACL (Supplementary Table 1). We required 95% of the read length to be aligned (-minp) ^70^. A hashsize (-hs) of 15 was specified. All hash positions were initially stored by the database, but only 100 random hash positions were kept for each seed (-mhp). In each seed cluster, we required a minimum length of 20 bp (-act). The bam-files were sorted using Samtools. Median coverage was calculated using BEDTools, ignoring positions that had not acquired any reads. To avoid mapping artefacts such as high coverage of only limited genomic regions, we required ≥40% breadth (fraction of genome covered by at least 1 read). We also required a median coverage depth of ≥20X to include a sample. Samples displaying coverage depth values higher than the threshold were downsampled to 20x using Samtools. To enable statistically sound population genomic comparisons along the transect and through time we required at least 5 samples to fulfill the coverage and breadth thresholds.

### SNP calling

SNP-calling was performed once per MAG, after combining BAM files from the approved samples into a multi-sample BAM file, using Freebayes ^69^, using the --pooled-continuous flag. SNPs were called only when supported by ≥4 reads and with an allele frequency of ≥1% ^71^. Also, we approved only bases with a Phred score >20 (corresponding to 99% probability of being a real SNP) to be included in downstream-analyses.

### POGENOM runs

POGENOM was run with the parameter settings --min_count 10, --subsample 10 and --min_found 1 on a VCF file of all approved samples for each MAG. With other words, it included for a sample only those loci with allele counts ≥10 (i.e. with ≥10 overlapping reads) and for those loci with counts >10, it downsampled to counts = 10. And overall, it included only those loci fulfilling the--min_count conditions for at least one sample. When analysing gene-specific *F*_ST_, --min_found was instead set to the same as the number of samples, i.e. restricting the calculations to loci with data for all samples. In this case, POGENOM was run on the same VCF files as before, but limiting the analysis to the surface transect samples by using the --sample_file flag. GFF files with gene definitions were obtained by using Prokka ^72^.

### Environmental association analysis

Nucleotide diversity was compared against temperature, salinity, chlorophyll *a*, DOC, TN, phosphate, nitrite, nitrate, ammonia and DON:DIN across the horizontal surface transect using Spearman’s rank correlation analysis in R ^73^.The p-values were False Discovery Rate adjusted. The same correlation analysis was conducted for nucleotide diversity over time (LMO) vs temperature, salinity, chlorophyll *a*, DOC and nitrate. The same environmental variables were included when the analysis was run collectively for surface and times series data (Surface+LMO). In order to study the potential association between diversity and time, we conducted multiple Mantel tests using distance matrices of π and days between sampling events. When diversity was studied across the vertical dimension we also included sampling depth and oxygen as independent variables.

For exploratory purposes, and in order to find the single best predictor of population genomic structure in individual cases, we conducted multiple pairwise tests between explanatory variables and genome-level *F*_ST_ values using the bioenv-function in R ^73^, initially allowing only one independent variable. Included environmental variables were the same as above, except that the spatial factor (MEM variables) was also taken into consideration. The statistical significance of these initial results were tested by Mantel’s tests, which uses the same definition of correlation as the bioenv-function. Prior to Mantel’s tests, the environmental variables were transformed into distance matrices and scaled to unit standard deviation. The p-values were False Discovery Rate adjusted and a *Q*-value <0.05 was considered statistically significant. Next, we repeated the bioenv-analysis but allowed three independent variables, in order to analyze the effect of multiple environmental variables on population structure. Multiple variables (with potential co-variation) were obtained in case the best model resulting from the bioenv-analysis contained more than one independent variable. However, such analyses may inflate Type-I error, but reduces the number of relevant explanatory variables which needs to be smaller than n-1 to avoid saturation of a Redundancy Analysis model (RDA) ^74^. Next, we conducted environmental association analyses in a global RDA analysis, followed by a conditioned analysis. This allowed us to disentangle the relative contribution of different explanatory variables in driving seascape genomic structure. The RDA was conducted using only the variables included in the most significant model suggested by the bioenv-function (three allowed). *F*_ST_ matrices were subjected to an unconstrained Principal coordinates analysis (PCoA) and the PC-axes were used as dependent input in the RDA. In the RDA, the regression coefficients are reported as adjusted values of multiple determination (*R^2^*-adj.). Statistical significance of the global RDA was evaluated using the permute-function from vegan and by performing an Anova (by “term”, 999 permutations) on the RDA to assess the statistical significance of each variable. The conditioned analysis was only conducted in case the global RDA showed statistically significant explanatory power (p < 0.05) to avoid Type I error and overestimation of the explained variance ^75^. Statistical significance of conditioned individual fractions (i.e. marginal effects) was evaluated by an Anova (by “margin”; 999 permutations).

## Declarations

### Ethics approval and consent to participate

Not applicable.

### Consent for publication

Not applicable.

### Availability of data and materials

The MAG sequences from Hugerth et al. ^34^ are available at NCBI’s Whole Genome Shotgun database under accession numbers LIAK00000000–LIDO00000000 and the LMO metagenome sequencing reads from the same study can be retrieved from NCBI’s Sequence Read Archive under accession numbers SRR2053273–SRR2053308. The preprocessed sequencing reads from the Transect samples from Alneberg et al. ^35^ are available at ENA hosted by EMBL-EBI under the study accession number PRJEB22997 (European Nucleotide Archive ERP104730). Source code and documentation for POGENOM are available at https://github.com/EnvGen/POGENOM.

### Authors’ contributions

AA initiated and led the project and coordinated and assisted with the data analysis. CS and AA wrote the manuscript. CS performed the bioinformatic analyses and environmental association analyses. AA wrote the POGENOM program. LF wrote the Snakemake workflow for automatic preparation of input data to POGENOM. JA assisted with bioinformatic analyses. All authors read and approved the final manuscript.

## Acknowledgements

We are grateful to Matthias Labrenz and Christin Bennke at IOW for early access to the Transect sample metagenomes. Computations were performed on resources provided by the Swedish National Infrastructure for Computing (SNIC) through the Uppsala Multidisciplinary Center for Advanced Computational Science (UPPMAX). We are grateful to Verena Kutschera at the National Bioinformatics Infrastructure Sweden (NBIS) for valuable input on the population genomics analysis.

## References

1. Azam, F. & Malfatti, F. Microbial structuring of marine ecosystems. Nat. Rev. Microbiol. 5, 782–791 (2007).

2. Dinsdale, E. A. et al. Functional metagenomic profiling of nine biomes. Nature 452, 629–632 (2008).

3. Giovannoni, S. J. & Stingl, U. Molecular diversity and ecology of microbial plankton. Nature vol. 437 343–348 (2005).

4. DeLong, E. F. et al. Community genomics among stratified microbial assemblages in the ocean’s interior. Science 311, 496–503 (2006).

5. Field, K. G. et al. Diversity and depth-specific distribution of SAR11 cluster rRNA genes from marine planktonic bacteria. Appl. Environ. Microbiol. 63, 63–70 (1997).

6. Morris, R. M. et al. SAR11 clade dominates ocean surface bacterioplankton communities. Nature 420, 806–810 (2002).

7. Dupont, C. L. et al. Functional tradeoffs underpin salinity-driven divergence in microbial community composition. PLoS One 9, e89549 (2014).

8. Jaspers, E. & Overmann, J. Ecological significance of microdiversity: identical 16S rRNA gene sequences can be found in bacteria with highly divergent genomes and ecophysiologies. Appl. Environ. Microbiol. 70, 4831–4839 (2004).

9. Konstantinidis, K. T. & DeLong, E. F. Genomic patterns of recombination, clonal divergence and environment in marine microbial populations. ISME J. 2, 1052–1065 (2008).

10. Nayfach, S., Rodriguez-Mueller, B., Garud, N. & Pollard, K. S. An integrated metagenomics pipeline for strain profiling reveals novel patterns of bacterial transmission and biogeography. Genome Res. 26, 1612–1625 (2016).

11. Hunt, D. E. et al. Resource partitioning and sympatric differentiation among closely related bacterioplankton. Science 320, 1081–1085 (2008).

12. Yoshida, M. et al. Intra-specific phenotypic and genotypic variation in toxic cyanobacterial Microcystis strains. J. Appl. Microbiol. 105, 407–415 (2008).

13. Johnson, Z. I. et al. Niche partitioning among Prochlorococcus ecotypes along ocean-scale environmental gradients. Science 311, 1737–1740 (2006).

14. Brown, M. V. et al. Global biogeography of SAR11 marine bacteria. Mol. Syst. Biol. 8, 595 (2012).

15. Giovannoni, S. J. et al. Genome streamlining in a cosmopolitan oceanic bacterium. Science 309, 1242–1245 (2005).

16. Carlson, C. A. et al. Seasonal dynamics of SAR11 populations in the euphotic and mesopelagic zones of the northwestern Sargasso Sea. ISME J. 3, 283–295 (2009).

17. Moore, L. R., Rocap, G. & Chisholm, S. W. Physiology and molecular phylogeny of coexisting Prochlorococcus ecotypes. Nature 393, 464–467 (1998).

18. Rodríguez-Valera, F. Approaches to prokaryotic biodiversity: a population genetics perspective. Environ. Microbiol. 4, 628–633 (2002).

19. Eloe-Fadrosh, E. A., Ivanova, N. N., Woyke, T. & Kyrpides, N. C. Metagenomics uncovers gaps in amplicon-based detection of microbial diversity. Nat Microbiol 1, 15032 (2016).

20. O’Brien, J. D. et al. A Bayesian approach to inferring the phylogenetic structure of communities from metagenomic data. Genetics 197, 925–937 (2014).

21. Luo, C. et al. ConStrains identifies microbial strains in metagenomic datasets. Nat. Biotechnol. 33, 1045–1052 (2015).

22. Quince, C. et al. DESMAN: a new tool for de novo extraction of strains from metagenomes. Genome Biol. 18, 181 (2017).

23. Schloissnig, S. et al. Genomic variation landscape of the human gut microbiome. Nature 493, 45–50 (2013).

24. Delmont, T. O. et al. Single-amino acid variants reveal evolutionary processes that shape the biogeography of a global SAR11 subclade. Elife 8, (2019).

25. Costea, P. I. et al. metaSNV: A tool for metagenomic strain level analysis. PLoS One 12, e0182392 (2017).

26. Gabrielsen, T. M., Brochmann, C. & Rueness, J. The Baltic Sea as a model system for studying postglacial colonization and ecological differentiation, exemplified by the red alga Ceramium tenuicorne. Mol. Ecol. 11, 2083–2095 (2002).

27. Johannesson, K. & André, C. INVITED REVIEW: Life on the margin: genetic isolation and diversity loss in a peripheral marine ecosystem, the Baltic Sea. Molecular Ecology vol. 15 2013–2029 (2006).

28. Ojaveer, H. et al. Status of biodiversity in the Baltic Sea. PLoS One 5, (2010).

29. Bergstrom, L., Tatarenkov, A., Johannesson, K., Jonsson, R. B. & Kautsky, L. GENETIC AND MORPHOLOGICAL IDENTIFICATION OF FUCUS RADICANS SP. NOV. (FUCALES, PHAEOPHYCEAE) IN THE BRACKISH BALTIC SEA1. Journal of Phycology vol. 41 1025–1038 (2005).

30. Jørgensen, H. B. H., Pertoldi, C., Hansen, M. M., Ruzzante, D. E. & Loeschcke, V. Genetic and environmental correlates of morphological variation in a marine fish: the case of Baltic Sea herring (Clupea harengus). Canadian Journal of Fisheries and Aquatic Sciences vol. 65 389–400 (2008).

31. Martinez Barrio, A. et al. The genetic basis for ecological adaptation of the Atlantic herring revealed by genome sequencing. Elife 5, (2016).

32. Sjöqvist, C., Godhe, A., Jonsson, P. R., Sundqvist, L. & Kremp, A. Local adaptation and oceanographic connectivity patterns explain genetic differentiation of a marine diatom across the North Sea-Baltic Sea salinity gradient. Mol. Ecol. 24, 2871–2885 (2015).

33. Herlemann, D. P. R. et al. Transitions in bacterial communities along the 2000 km salinity gradient of the Baltic Sea. The ISME Journal vol. 5 1571–1579 (2011).

34. Hugerth, L. W. et al. Metagenome-assembled genomes uncover a global brackish microbiome. Genome Biol. 16, 279 (2015).

35. Alneberg, J. et al. BARM and BalticMicrobeDB, a reference metagenome and interface to meta-omic data for the Baltic Sea. Scientific Data vol. 5 (2018).

36. Alneberg, J. et al. Genomes from uncultivated prokaryotes: a comparison of metagenome-assembled and single-amplified genomes. Microbiome 6, 173 (2018).

37. Alneberg, J. et al. Binning metagenomic contigs by coverage and composition. Nature Methods vol. 11 1144–1146 (2014).

38. Mcdonald, J. H. & Kreitman, M. Neutral mutation hypothesis test. Nature vol. 354 116–116 (1991).

39. Oren, A. Bioenergetic aspects of halophilism. Microbiol. Mol. Biol. Rev. 63, 334–348 (1999).

40. Gevers, D. et al. Opinion: Re-evaluating prokaryotic species. Nat. Rev. Microbiol. 3, 733–739 (2005).

41. Herlemann, D. P. R., Lundin, D., Andersson, A. F., Labrenz, M. & Jürgens, K. Phylogenetic Signals of Salinity and Season in Bacterial Community Composition Across the Salinity Gradient of the Baltic Sea. Front. Microbiol. 7, 1883 (2016).

42. Schattenhofer, M. et al. Latitudinal distribution of prokaryotic picoplankton populations in the Atlantic Ocean. Environ. Microbiol. 11, 2078–2093 (2009).

43. Yooseph, S. et al. Genomic and functional adaptation in surface ocean planktonic prokaryotes. Nature 468, 60–66 (2010).

44. Treusch, A. H. et al. Seasonality and vertical structure of microbial communities in an ocean gyre. The ISME Journal vol. 3 1148–1163 (2009).

45. Excoffier, L. & Ray, N. Surfing during population expansions promotes genetic revolutions and structuration. Trends Ecol. Evol. 23, 347–351 (2008).

46. Wright, S. Evolution in Mendelian Populations. Genetics 16, 97–159 (1931).

47. Nei, M. Analysis of Gene Diversity in Subdivided Populations. Proceedings of the National Academy of Sciences vol. 70 3321–3323 (1973).

48. Shapiro, B. J. et al. Population genomics of early events in the ecological differentiation of bacteria. Science 336, 48–51 (2012).

49. Charlesworth, B. Effective population size and patterns of molecular evolution and variation. Nature Reviews Genetics vol. 10 195–205 (2009).

50. Berg, P. R. et al. Adaptation to Low Salinity Promotes Genomic Divergence in Atlantic Cod (Gadus morhua L.). Genome Biology and Evolution vol. 7 1644–1663 (2015).

51. Feder, J. L., Egan, S. P. & Nosil, P. The genomics of speciation-with-gene-flow. Trends in Genetics vol. 28 342–350 (2012).

52. Fernández-Gómez, B. et al. Patterns and architecture of genomic islands in marine bacteria. BMC Genomics 13, 347 (2012).

53. Qin, Q.-L. et al. Comparative genomics of the marine bacterial genusGlaciecolareveals the high degree of genomic diversity and genomic characteristic for cold adaptation. Environmental Microbiology vol. 16 1642–1653 (2014).

54. Hahn, M. W. et al. Isolation of novel ultramicrobacteria classified as actinobacteria from five freshwater habitats in Europe and Asia. Appl. Environ. Microbiol. 69, 1442–1451 (2003).

55. Glöckner, F. O. et al. Comparative 16S rRNA analysis of lake bacterioplankton reveals globally distributed phylogenetic clusters including an abundant group of actinobacteria. Appl. Environ. Microbiol. 66, 5053–5065 (2000).

56. Neuenschwander, S. M., Ghai, R., Pernthaler, J. & Salcher, M. M. Microdiversification in genome-streamlined ubiquitous freshwater Actinobacteria. ISME J. 12, 185–198 (2018).

57. Li, W. K. W. Annual average abundance of heterotrophic bacteria and Synechococcus in surface ocean waters. Limnology and Oceanography vol. 43 1746–1753 (1998).

58. Haverkamp, T. H. A. et al. Colorful microdiversity of Synechococcus strains (picocyanobacteria) isolated from the Baltic Sea. ISME J. 3, 397–408 (2009).

59. Szabo, G. et al. Reproducibility of Vibrionaceae population structure in coastal bacterioplankton. ISME J. 7, 509–519 (2013).

60. Coleman, M. L. et al. Genomic islands and the ecology and evolution of Prochlorococcus. Science 311, 1768–1770 (2006).

61. Kettler, G. C. et al. Patterns and implications of gene gain and loss in the evolution of Prochlorococcus. PLoS Genet. 3, e231 (2007).

62. Laikre, L., Palm, S. & Ryman, N. Genetic population structure of fishes: implications for coastal zone management. Ambio 34, 111–119 (2005).

63. Cabello-Yeves, P. J. & Rodriguez-Valera, F. Marine-freshwater prokaryotic transitions require extensive changes in the predicted proteome. Microbiome 7, 117 (2019).

64. Kelly, R. P. & Palumbi, S. R. Genetic structure among 50 species of the northeastern Pacific rocky intertidal community. PLoS One 5, e8594 (2010).

65. Langmead, B. & Salzberg, S. L. Fast gapped-read alignment with Bowtie 2. Nat. Methods 9, 357–359 (2012).

66. Li, H. & Durbin, R. Fast and accurate short read alignment with Burrows-Wheeler transform. Bioinformatics vol. 25 1754–1760 (2009).

67. Lee, W.-P. et al. MOSAIK: a hash-based algorithm for accurate next-generation sequencing short-read mapping. PLoS One 9, e90581 (2014).

68. Van der Auwera, G. A. et al. From FastQ data to high confidence variant calls: the Genome Analysis Toolkit best practices pipeline. Curr. Protoc. Bioinformatics 43, 11.10.1–11.10.33 (2013).

69. Garrison E. & Marth G. Haplotype-based variant detection from short-read sequencing. arXiv preprint arXiv:1207.3907 [q-bio.GN] (2012).

70. Konstantinidis, K. T. & Tiedje, J. M. Prokaryotic taxonomy and phylogeny in the genomic era: advancements and challenges ahead. Current Opinion in Microbiology vol. 10 504–509 (2007).

71. 1000 Genomes Project Consortium et al. A map of human genome variation from population-scale sequencing. Nature 467, 1061–1073 (2010).

72. Seemann, T. Prokka: rapid prokaryotic genome annotation. Bioinformatics 30, 2068–2069 (2014).

73. R Core Team (2017). R: A language and environment for statistical computing. R Foundation for Statistical Computing, Vienna, Austria URL https://www.R-project.org/.

74. Borcard, D., Gillet, F. & Legendre, P. Numerical Ecology with R. (2011) doi:10.1007/978-1-4419-7976-6.

75. Blanchet, F. G., Legendre, P. & Borcard, D. Forward selection of explanatory variables. Ecology 89, 2623–2632 (2008).

