## Supplementary Information for "Revealing ecologically coherent population structure of uncultivated bacterioplankton with POGENOM"

**Table of contents**


1. Calculation of population genetic parameters in POGENOM

    1.1. Nucleotide diversity, $\pi$

    1.2. Fixation index, $F_{ST}$

    1.3. Amino-acid level $\pi$ and $F_{ST}$

    1.4. *pN/pS*

2. Supplementary figures

3. Supplementary tables

### 1. Calculation of population genetic parameters in POGENOM

#### 1.1. Nucleotide diversity, $\pi$

Nucleotide diversity ($\pi$), sometimes called heterozygosity, is defined as the average number of nucleotide differences per site between any two sequence reads chosen randomly from the sample population ($0 \leq \pi < 1$). POGENOM calculates $\pi$ of a single locus according to Schloissnig et al. [23]:

$$\pi_i = \sum_{B_1 \in \{ACTG\}} \sum_{B_2 \in \{ACTG\} \setminus B_1} \frac{x_{i,B_1 S_1}}{c_i} \frac{x_{i,B_2}}{c_i - 1}$$

where $x_{i,Bj}$ is the counts of nucleotide $B_j$ at position $i$ in the genome for the sample, and $c_i$ is the total coverage (sequence depth) at position $i$ for the sample. To calculate a genome-wide $\pi$, $\pi$ is averaged over all loci by summing all $\pi_i$ and dividing by the genome size. Loci not included in the VCF file are assumed to lack diversity (to have $\pi_i = 0$). To calculate a gene-wise $\pi$, $\pi$ is instead averaged over all loci within the gene, including the start codon but excluding the stop codon. The $\pi$ calculation also works for alleles >1 bp (in case a variant caller was used that output haplotypes), then instead basing the calculations on counts of haplotypes (oligomers) rather than nucleotides for loci where alleles >1 bp are reported in the VCF file. POGENOM can also split counts of haplotypes into counts of individual nucleotides, if this is preferred (this is the default). For gene-wise $\pi$, $\pi$ will for a gene and a sample per default be set to NA if one or several loci in the gene included in the VCF file have missing data for the sample, to avoid biases between samples for the gene due to missing data.

A locus not reported in the VCF file can be missing because no genetic variation was observed and/or because the locus did not have sufficient sequence depth coverage for the pool of samples when running the variant calling. The latter can lead to $\pi$ values being biased downwards (since these loci are assumed to have $\pi_i = 0$) and can skew the comparison of $\pi$ between genomes (but should however not affect comparisons between samples for the same genome, if multi-sample variant calling was conducted). In order to adjust for this, a normalised genome-wide $\pi$ is also calculated for each sample by dividing the genome-wide $\pi$ with an estimated completeness factor for the sample. The completeness factor is based on the assumption that loci with sufficient coverage (exceeding the cutoff applied in POGENOM) in one sample are independent from those in another sample. Thus, the loci covered in sample 1 can be treated as a random subset of the genome and used to assess the completeness of sample 2, by calculating what fraction of loci covered by sample 1 are also covered by sample 2. The completeness for a sample is assessed this way using all other samples, and the completeness factor is the average of these assessments. The normalised genome-wide $\pi$ is not used for $F_{ST}$ calculations since missing variant loci should affect intra- and intersample $\pi$ equally and have little influence on the $F_{ST}$ (see calculations below).

#### 1.2. Fixation index, $F_{ST}$

To calculate the fixation index ($F_{ST}$) for each pair of samples, the intersample $\pi$ has to be calculated. For a single locus this is calculated according to[23]:

$$\pi_{i,S_1,S_2} = \sum_{B_1 \in \{ACTG\}} \sum_{B_2 \in \{ACTG\} \setminus B_1} \frac{x_{i,B_1,S_1}}{c_{i,S_1}} \frac{x_{i,B_2,S_2}}{c_{i,S_2}}$$

where $x_{i,Bj,Sk}$ is the number of nucleotide $B_j$ observed at position *i* in the genome in sample $S_k$ and $c_{i,Sk}$ is the coverage of position *i* in sample $S_k$. The inter-sample $\pi$ is then calculated for the whole genome (or gene) by summing $\pi_i$ for all loci (or loci inside the gene) and dividing by the genome (or gene) size.

$F_{ST}$ is then calculated according to:

$$F_{ST} = 1 - \frac{mean\ (\pi_{intra\ sample})}{\pi_{inter\ sample}} = 1 - \frac{(\pi_{S_1} + \pi_{S_2})/2}{\pi_{S_1,S_2}}$$

where for genome-wide $F_{ST}$, the calculation is based on genome-wide intra- and intersample $\pi$ values, while for gene-wise $F_{ST}$ it is based on gene-wise $\pi$ values. For both types of $F_{ST}$ calculations, only loci for which both samples in the pair have data will be considered for the intra- and intersample $\pi$ calculations. If no such loci are present, or if the intersample $\pi$ is zero (which only happens if also both of the intrasample $\pi$ are zero), $F_{ST}$ will be set to NA.

For permuted gene-wise $F_{ST}$, the variant loci are randomly redistributed among the genes in a way that each gene will obtain a new set of variant loci (with their associated allele frequencies) but will have the same number of variant loci as in the original case.

### 1.3. Amino-acid level $\pi$ and $F_{ST}$

Gene-wise amino acid $\pi$ is calculated based on the variant loci within genes, including the start codon but excluding the stop codon. Amino acid $\pi$ for a single locus is calculated by modifying the gene sequence according to each detected allele (one at a time) for the locus in the sample and translating the modified gene into a peptide (based on the genetic code file). The counts of each unique peptide will then be used for the calculations of intra- and intersample $\pi$ (rather than the counts of individual nucleotides [or haplotypes] as above). This approach allows adequate amino acid diversity calculations also when having alleles >1 bp (haplotypes). The gene-wise amino acid level $F_{ST}$ is calculated analogously to the gene-wise nucleotide level $F_{ST}$ from the amino acid intra- and intersample $\pi$ values. As for gene-wise nucleotide diversity, a gene will for a sample get $\pi$ = NA if one or several loci in the gene that are included in the VCF file have missing data for the sample.

## 1.4. *pN/pS*

*pN/pS* measures the ratio of the nonsynonymous to the synonymous polymorphism rates, where *pN* equals the fraction of possible nonsynonymous mutations that are observed as polymorphisms and *pS* equals the fraction of synonymous mutations that are observed as polymorphisms. To calculate the *pN/pS* for a gene and sample, POGENOM first derives a consensus nucleotide sequence for the gene in the sample by modifying the reference nucleotide in variant loci based on the most frequent allele, while keeping the reference sequence in invariant positions. For each nucleotide position, every possible (single nucleotide) mutation relative to the consensus sequence is then recorded, and whether this mutation is nonsynonymous or synonymous and present as a polymorphism or not.

*pN*/*pS* is then calculated as:

$$pN/pS = \frac{\left[\frac{\sum_{i=1}^{L} n_i}{\sum_{i=1}^{L} N_i}\right]}{\left[\frac{\sum_{i=1}^{L} s_i}{\sum_{i=1}^{L} S_i}\right]}$$

where $n_i$ is the number of observed nonsynonymous mutations (alleles), $N_i$ is the total number of possible nonsynonymous mutations, $s_i$ is the number of observed synonymous mutations and $S_i$ is the total number of possible nonsynonymous mutations for locus *i.* If no synonymous mutations are observed for the gene *pN/pS* is set to NA. In addition to calculating *pN/pS* on a per-sample basis, POGENOM calculates it on all samples collectively by combing the allele frequencies of all samples.

### 2. Supplementary figures

**Supplementary Figure 1**. (**a**) Mean nucleotide diversity of single-copy core genes (SCG) compared against all other genes in the respective BACL. (**b**) Mean pN/pS of SCGs vs. other genes per BACL.

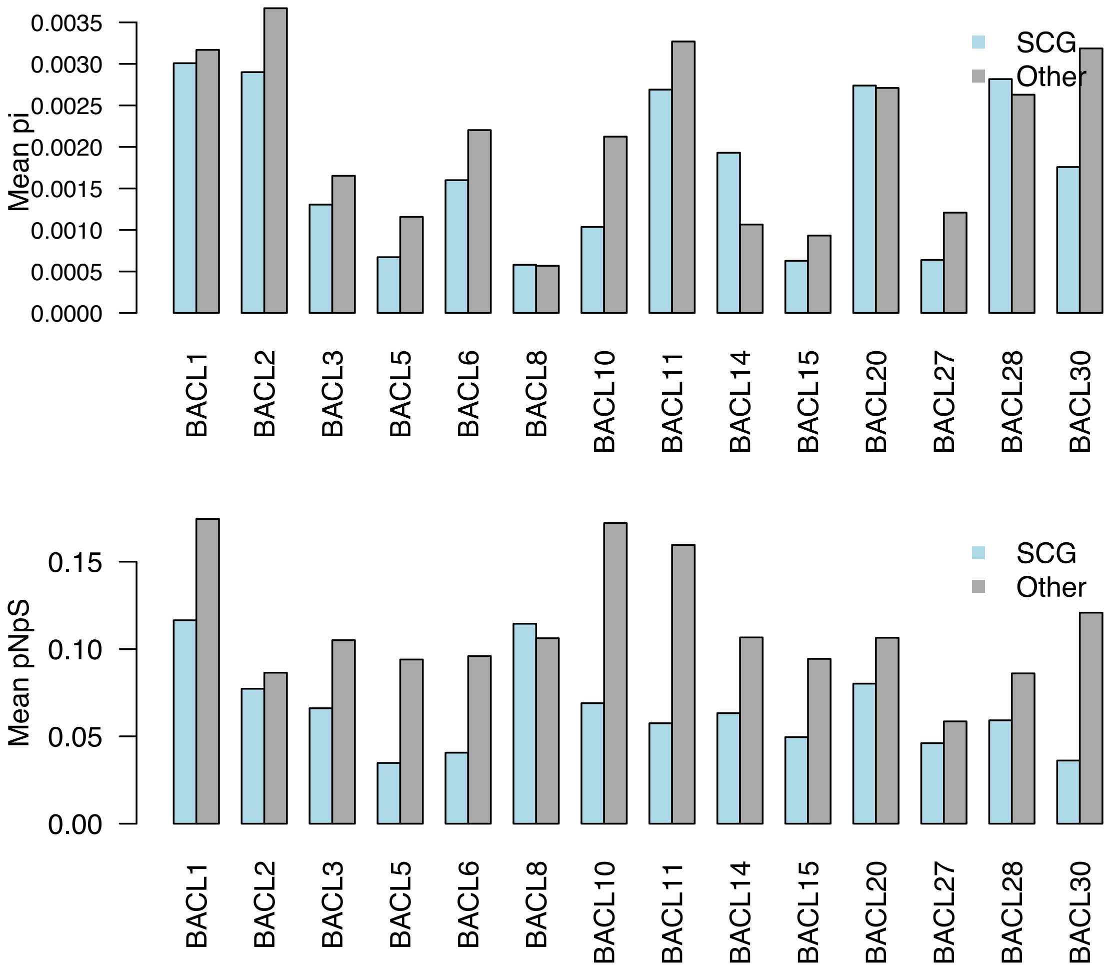

**Supplementary Figure 2**. Variation in proportion of acid (D (aspartic acid), E (glutamic acid)) and alkalic (H (histidine), K (lysine), R (arginine)) amino acids along the salinity gradient for different BACLs. Each bar represents one sample. The numbers in the top indicate the salinity values of the samples, from left to right.

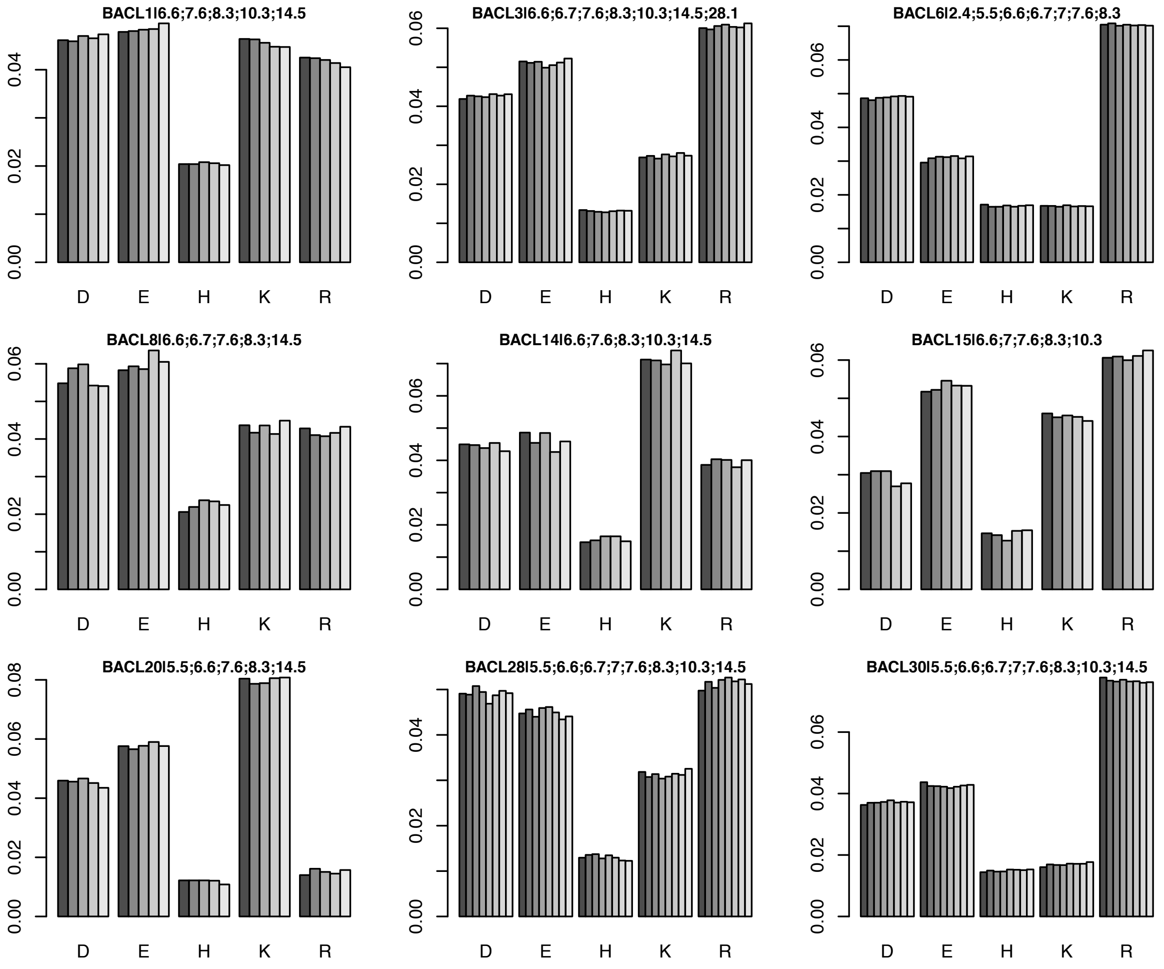

**Supplementary Figure 3**. Isolation by distance (IBD) for pairwise comparisons of all Transect samples. Genetic differentiation measured as $F_{ST}$. Geographical distance between respective sample sites as nautical miles (nm).

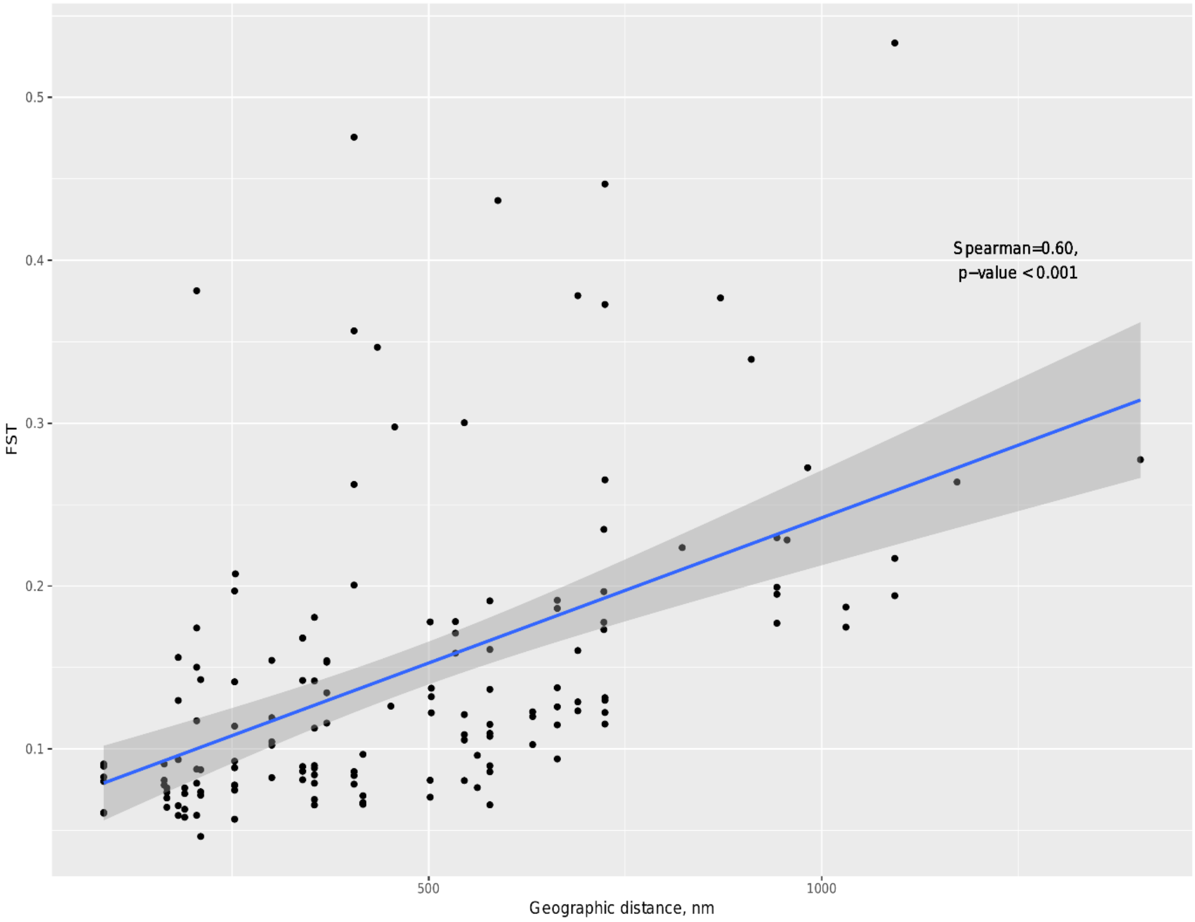

### 3. Supplementary tables

**Supplementary Table 1**. Summary statistics of MAG clusters (Baltic Clusters; BACL) produced in Hugerth et al (2015). One MAG (highest N50 length) from each cluster was chosen as a reference genome for the population genomic analysis. Number of loci only applicable for BACL included in downstream analysis.

| Baltic Cluster | MAG ID | Number of contigs | Number of bases | N50 | N50 length | N90 | N90 length | Longest contig | Coverage within sample | Number of loci | Phylogeny |
|---|---|---|---|---|---|---|---|---|---|---|---|
| BACL1 | 120820_bin45 | 213 | 1455539 | 25 | 16167 | 118 | 2482 | 101813 | 37 | 29457 | BACTERIA;PROTEOBACTERIA;GAMMAPROTEOBACTERIA;SAR86; |
| BACL2 | 120802_bin41 | 311 | 1101056 | 64 | 5183 | 223 | 1510 | 25328 | 23 | 15294 | BACTERIA;ACTINOBACTERIA;ACI; |
| BACL3 | 120619_bin3 | 474 | 2648901 | 70 | 10134 | 290 | 2185 | 66229 | 20 | 15357 | BACTERIA;PROTEOBACTERIA;GAMMAPROTEOBACTERIA;OM182; |
| BACL4 | 120920_bin74 | 259 | 1080997 | 40 | 7155 | 174 | 1555 | 54245 | 24 | NA | BACTERIA;ACTINOBACTERIA;ACI; |
| BACL5 | 120820_bin39 | 228 | 1151850 | 38 | 8535 | 149 | 2067 | 39080 | 32 | 3686 | BACTERIA;PROTEOBACTERIA;ALPHAPROTEOBACTERIA;SAR11; |
| BACL6 | 120910_bin40 | 269 | 1688323 | 51 | 10573 | 172 | 2699 | 35216 | 17 | 12406 | BACTERIA;ACTINOBACTERIA;ACIV; |
| BACL7 | 120322_bin74 | 77 | 1743356 | 12 | 49392 | 42 | 11419 | 136363 | 27 | NA | BACTERIA;BACTEROIDETES;FLAVOBACTERIIA;CRYOMORPHACEAE; |
| BACL8 | 120531_bin13 | 208 | 2086531 | 24 | 22088 | 109 | 4301 | 120247 | 37 | 3679 | BACTERIA;BACTEROIDETES;FLAVOBACTERIIA;FLAVOBACTERIACEAE; |
| BACL9 | 120507_bin52 | 458 | 1601779 | 92 | 5174 | 328 | 1545 | 40179 | 21 | NA | BACTERIA;VERRUCOMICROBIA;LD19; |
| BACL10 | 120910_bin24 | 671 | 2763624 | 116 | 6842 | 452 | 1571 | 66984 | 23 | 23939 | BACTERIA;PROTEOBACTERIA;ALPHAPROTEOBACTERIA;RHODOBACTER; |
| BACL11 | 121015_bin20 | 475 | 1209326 | 124 | 2935 | 373 | 1341 | 16423 | 18 | 13090 | BACTERIA;BACTEROIDETES;FLAVOBACTERIIA;CRYOMORPHACEAE; |
| BACL12 | 120813_bin55 | 721 | 2799617 | 156 | 5133 | 521 | 1766 | 32481 | 14 | NA | BACTERIA;BACTEROIDETES;SPHINGOBACTERIIA;SPHINGOBACTERIALES; |
| BACL13 | 121220_bin23 | 183 | 1270387 | 30 | 13618 | 111 | 2708 | 67392 | 26 | NA | ARCHAEA;THAUMARCHAEOTA;NITROSOPUMILUS; |
| BACL14 | 120920_bin58 | 268 | 1280196 | 46 | 8793 | 168 | 1664 | 29068 | 21 | 4032 | BACTERIA;PROTEOBACTERIA;BETAPROTEOBACTERIA;OM43; |
| BACL15 | 120619_bin91 | 243 | 1214420 | 40 | 8106 | 161 | 2161 | 36531 | 35 | 3353 | BACTERIA;ACTINOBACTERIA;ACI; |
| BACL16 | 120619_bin48 | 101 | 2527476 | 13 | 55966 | 47 | 11566 | 181472 | 27 | NA | BACTERIA;PROTEOBACTERIA;GAMMAPROTEOBACTERIA;SAR92; |
| BACL17 | 120823_bin42 | 255 | 1352430 | 53 | 8060 | 175 | 2443 | 32953 | 23 | NA | BACTERIA;ACTINOBACTERIA;ACIV; |
| BACL18 | 120924_bin36 | 455 | 1459716 | 106 | 4033 | 339 | 1526 | 19136 | 23 | NA | BACTERIA;BACTEROIDETES;FLAVOBACTERIIA;CRYOMORPHACEAE; |
| BACL19 | 120924_bin39 | 496 | 1258466 | 124 | 2981 | 386 | 1312 | 21420 | 20 | NA | BACTERIA;ACTINOBACTERIA;ACIV; |
| BACL20 | 120920_bin64 | 510 | 1143837 | 135 | 2586 | 407 | 1186 | 12816 | 44 | 10510 | BACTERIA;PROTEOBACTERIA;ALPHAPROTEOBACTERIA;SAR11; |
| BACL21 | 121220_bin10 | 109 | 1915951 | 21 | 28212 | 64 | 10000 | 88771 | 22 | NA | BACTERIA;BACTEROIDETES;FLAVOBACTERIIA;FLAVOBACTERIACEAE; |
| BACL22 | 120619_bin32 | 520 | 2408986 | 103 | 6612 | 359 | 2177 | 27387 | 14 | NA | BACTERIA;BACTEROIDETES;FLAVOBACTERIIA;FLAVOBACTERIACEAE; |
| BACL23 | 120924_bin60 | 324 | 1727489 | 67 | 7321 | 219 | 2483 | 36814 | 39 | NA | BACTERIA;BACTEROIDETES;FLAVOBACTERIIA;CRYOMORPHACEAE; |
| BACL24 | 120322_bin51 | 567 | 2980252 | 66 | 11906 | 339 | 1745 | 57853 | 29 | NA | BACTERIA;VERRUCOMICROBIA;OPITUTACEAE; |
| BACL25 | 120322_bin65 | 160 | 1262816 | 35 | 10273 | 116 | 3731 | 44217 | 19 | NA | BACTERIA;ACTINOBACTERIA;LUNA; |
| BACL26 | 121220_bin70 | 743 | 1907425 | 199 | 2986 | 580 | 1359 | 20803 | 17 | NA | BACTERIA;PROTEOBACTERIA;GAMMAPROTEOBACTERIA;SAR92; |
| BACL27 | 120823_bin4 | 658 | 1672064 | 152 | 2954 | 510 | 1263 | 27311 | 37 | 5812 | BACTERIA;ACTINOBACTERIA;ACIV; |
| BACL28 | 120531_bin53 | 307 | 1121591 | 72 | 4744 | 224 | 1688 | 19072 | 15 | 9790 | BACTERIA;ACTINOBACTERIA;LUNA; |
| BACL29 | 121220_bin8 | 518 | 1480518 | 136 | 3426 | 397 | 1466 | 14929 | 11 | NA | BACTERIA;BACTEROIDETES;FLAVOBACTERIIA;FLAVOBACTERIACEAE; |
| BACL30 | 120619_bin27 | 570 | 1813814 | 130 | 4363 | 421 | 1405 | 20340 | 24 | 21122 | BACTERIA;CYANOBACTERIA;CYANOBIUM; |

**Supplementary Table 2.** Information about sampling stations, sampling depths and dates. Abbreviation column, s=surface, m=mid, d=deep.

| Station | Abrreviation | Depth, m | Latitude | Longitude | Sampling date |
|---|---|---|---|---|---|
| Bothnian Bay | BB_s | 2 | 65.45 | 23.30 | 2014-06-12 |
| Bothnian Bay | BB_m | 43 | 65.45 | 23.30 | 2014-06-12 |
| Bothnian Bay | BB_d | 79 | 65.45 | 23.30 | 2014-06-12 |
| Bothnian Bay | BS_s | 2 | 61.78 | 19.30 | 2014-06-11 |
| Bothnian Bay | BS_m | 34 | 61.78 | 19.30 | 2014-06-11 |
| Bothnian Bay | BS_d | 56 | 61.78 | 19.30 | 2014-06-11 |
| Baltic Proper | BP_N_s | 2 | 58.58 | 18.23 | 2014-06-15 |
| Baltic Proper | BP_N_m | 61 | 58.58 | 18.23 | 2014-06-15 |
| Baltic Proper | BP_N_d | 76 | 58.58 | 18.23 | 2014-06-15 |
| Baltic Proper | BP_E_s | 2 | 57.31 | 20.08 | 2014-06-09 |
| Baltic Proper | BP_E_m | 65 | 57.31 | 20.08 | 2014-06-09 |
| Baltic Proper | BP_E_d | 116 | 57.31 | 20.08 | 2014-06-09 |
| Baltic Proper | BP_W_s | 2 | 57.12 | 17.66 | 2014-06-17 |
| Baltic Proper | BP_W_m | 56 | 57.12 | 17.66 | 2014-06-17 |
| Baltic Proper | BP_W_d | 86 | 57.12 | 17.66 | 2014-06-17 |
| Baltic Proper | BP_S_s | 2 | 55.57 | 16.37 | 2014-06-08 |
| Baltic Proper | BP_S_m | 30 | 55.57 | 16.37 | 2014-06-08 |
| Baltic Proper | BP_S_d | 67 | 55.57 | 16.37 | 2014-06-08 |
| Arkona (North) | Arkona_N_s | 5 | 54.70 | 12.70 | 2014-06-04 |
| Arkona (North) | Arkona_N_m | 10 | 54.70 | 12.70 | 2014-06-04 |
| Arkona (North) | Arkona_N_d | 15 | 54.70 | 12.70 | 2014-06-04 |
| Arkona (South) | Arkona_S_s | 2 | 54.28 | 11.57 | 2014-06-07 |
| Arkona (South) | Arkona_S_m | 16 | 54.28 | 11.57 | 2014-06-07 |
| Arkona (South) | Arkona_S_d | 23 | 54.28 | 11.57 | 2014-06-07 |
| Öresund | Oresund_s | 2 | 56.00 | 11.08 | 2014-06-06 |
| Öresund | Oresund_m | 11 | 56.00 | 11.08 | 2014-06-06 |
| Öresund | Oresund_d | 21 | 56.00 | 11.08 | 2014-06-06 |
| Skagerrak | Skagerrak_s | 2 | 58.13 | 9.99 | 2014-06-05 |
| Skagerrak | Skagerrak_m | 80 | 58.13 | 9.99 | 2014-06-05 |
| Skagerrak | Skagerrak_d | 242 | 58.13 | 9.99 | 2014-06-05 |
| LMO | 120314 | 2 | 56.55 | 17.03 | 2012-03-12 |
| LMO | 120328 | 2 | 56.55 | 17.03 | 2012-03-28 |
| LMO | 120403 | 2 | 56.55 | 17.03 | 2012-04-03 |
| LMO | 120416 | 2 | 56.55 | 17.03 | 2012-04-16 |
| LMO | 120516 | 2 | 56.55 | 17.03 | 2012-05-16 |
| LMO | 120531 | 2 | 56.55 | 17.03 | 2012-05-31 |
| LMO | 120604 | 2 | 56.55 | 17.03 | 2012-06-04 |
| LMO | 120613 | 2 | 56.55 | 17.03 | 2012-06-13 |
| LMO | 120619 | 2 | 56.55 | 17.03 | 2012-06-19 |
| LMO | 120628 | 2 | 56.55 | 17.03 | 2012-06-28 |
| LMO | 120705 | 2 | 56.55 | 17.03 | 2012-07-05 |
| LMO | 120709 | 2 | 56.55 | 17.03 | 2012-07-09 |
| LMO | 120717 | 2 | 56.55 | 17.03 | 2012-07-17 |
| LMO | 120802 | 2 | 56.55 | 17.03 | 2012-08-02 |
| LMO | 120806 | 2 | 56.55 | 17.03 | 2012-08-06 |
| LMO | 120813 | 2 | 56.55 | 17.03 | 2012-08-13 |
| LMO | 120820 | 2 | 56.55 | 17.03 | 2012-08-20 |
| LMO | 120823 | 2 | 56.55 | 17.03 | 2012-08-23 |
| LMO | 120828 | 2 | 56.55 | 17.03 | 2012-08-28 |
| LMO | 120903 | 2 | 56.55 | 17.03 | 2012-09-03 |
| LMO | 120920 | 2 | 56.55 | 17.03 | 2012-09-20 |
| LMO | 120924 | 2 | 56.55 | 17.03 | 2012-09-24 |
| LMO | 121001 | 2 | 56.55 | 17.03 | 2012-10-01 |
| LMO | 121004 | 2 | 56.55 | 17.03 | 2012-10-04 |
| LMO | 121015 | 2 | 56.55 | 17.03 | 2012-10-15 |
| LMO | 121022 | 2 | 56.55 | 17.03 | 2012-10-22 |
| LMO | 121028 | 2 | 56.55 | 17.03 | 2012-10-28 |
| LMO | 121105 | 2 | 56.55 | 17.03 | 2012-11-05 |
| LMO | 121128 | 2 | 56.55 | 17.03 | 2012-11-28 |
| LMO | 121220 | 2 | 56.55 | 17.03 | 2012-12-20 |

**Supplementary Table 3**. Median coverage per sample and genome. Values equal to or greater than 20X as bold.

| Sample | BACL1 | BACL2 | BACL3 | BACL4 | BACL5 | BACL6 | BACL7 | BACL8 | BACL9 | BACL10 | BACL11 | BACL12 | BACL13 | BACL14 | BACL15 | BACL16 | BACL17 | BACL18 | BACL19 | BACL20 | BACL21 | BACL22 | BACL23 | BACL24 | BACL25 | BACL26 | BACL27 | BACL28 | BACL29 | BACL30 |
|---|---|---|---|---|---|---|---|---|---|---|---|---|---|---|---|---|---|---|---|---|---|---|---|---|---|---|---|---|---|---|
| BB_s | 4 | 88 | 1 | 6 | 1 | 47 | 5 | 1 | 3 | 12 | 1 | 2 | 2 | 10 | 5 | 1 | 4 | 8 | 4 | 1 | 2 | 2 | 1 | 2 | 3 | 1 | 23 | 2 | 1 | 10 |
| BB_m | 5 | 141 | 1 | 17 | 1 | 126 | 42 | 2 | 6 | 33 | 5 | 2 | 37 | 2 | 1 | 1 | 2 | 2 | 13 | 14 | 8 | 3 | 2 | 2 | 2 | 1 | 47 | 1 | 2 | 3 |
| BB_d | 1 | 84 | 2 | 20 | 2 | 98 | 60 | 1 | 6 | 33 | 3 | 2 | 17 | 2 | 3 | 3 | 4 | 5 | 7 | 3 | 9 | 1 | 69 | 6 | 3 | 2 | 65 | 1 | 2 | 8 |
| BS_s | 11 | 43 | 6 | 8 | 3 | 99 | 14 | 5 | 1 | 16 | 2 | 1 | 1 | 4 | 8 | 7 | 3 | 6 | 33 | 22 | 6 | 4 | 1 | 3 | 2 | 3 | 77 | 34 | 1 | 418 |
| BS_m | 43 | 130 | 4 | 24 | 29 | 335 | 24 | 4 | 2 | 51 | 6 | 1 | 5 | 15 | 23 | 2 | 12 | 4 | 28 | 2 | 18 | 21 | 1 | 10 | 4 | 2 | 155 | 32 | 1 | 119 |
| BS_d | 135 | 276 | 2 | 29 | 29 | 95 | 90 | 3 | 12 | 84 | 43 | 2 | 134 | 10 | 10 | 12 | 1 | 6 | 11 | 131 | 55 | 14 | 2 | 9 | 6 | 34 | 35 | 11 | 12 | 12 |
| BP_N_s | 56 | 115 | 39 | 17 | 11 | 56 | 44 | 74 | 3 | 18 | 2 | 1 | 4 | 31 | 94 | 7 | 2 | 2 | 3 | 108 | 31 | 15 | 2 | 1 | 1 | 1 | 10 | 210 | 1 | 323 |
| BP_N_m | 4 | 1 | 2 | 1 | 2 | 986 | 2 | 1 | 1 | 60 | 3 | 1 | 2 | 7 | 4 | 2 | 2 | 2 | 1 | 19 | 4 | 2 | 1 | 1 | 343 | 1 | 3 | 2 | 2 | 10 |
| BP_N_d | 6 | 2 | 1 | 1 | 4 | 264 | 18 | 3 | 15 | 41 | 4 | 2 | 66 | 2 | 1 | 8 | 1 | 1 | 2 | 2 | 10 | 2 | 1 | 2 | 24 | 4 | 1 | 2 | 3 | 2 |
| BP_E_s | 1 | 5 | 21 | 7 | 2 | 60 | 12 | 76 | 2 | 16 | 1 | 1 | 1 | 7 | 4 | 40 | 21 | 8 | 5 | 2 | 37 | 5 | 2 | 1 | 125 | 1 | 1 | 102 | 1 | 1619 |
| BP_E_m | 2 | 1 | 2 | 1 | 2 | 41 | 65 | 1 | 2 | 27 | 2 | 3 | 2 | 2 | 2 | 14 | 2 | 2 | 1 | 2 | 53 | 2 | 24 | 1 | 202 | 17 | 1 | 1 | 1 | 5 |
| BP_E_d | 14 | 2 | 1 | 1 | 6 | 110 | 4 | 1 | 5 | 15 | 1 | 1 | 2 | 4 | 1 | 2 | 1 | 2 | 1 | 1 | 4 | 3 | 4 | 1 | 16 | 11 | 4 | 1 | 2 | 9 |
| BP_W_s | 1 | 5 | 7 | 1 | 2 | 73 | 7 | 7 | 1 | 9 | 1 | 1 | 3 | 4 | 32 | 5 | 4 | 26 | 2 | 3 | 2 | 3 | 1 | 1 | 2 | 2 | 3 | 179 | 2 | 706 |
| BP_W_m | 5 | 2 | 2 | 2 | 2 | 497 | 26 | 1 | 8 | 48 | 20 | 2 | 48 | 6 | 1 | 5 | 3 | 2 | 4 | 10 | 8 | 5 | 1 | 2 | 90 | 25 | 2 | 1 | 1 | 17 |
| BP_W_d | 16 | 1 | 1 | 2 | 2 | 636 | 67 | 6 | 3 | 99 | 21 | 2 | 58 | 1 | 2 | 6 | 1 | 2 | 1 | 6 | 13 | 3 | 1 | 4 | 115 | 40 | 4 | 2 | 6 | 4 |
| BP_S_s | 69 | 17 | 103 | 4 | 4 | 87 | 23 | 95 | 4 | 27 | 1 | 41 | 2 | 103 | 73 | 10 | 6 | 10 | 1 | 211 | 10 | 9 | 6 | 1 | 2 | 2 | 2 | 203 | 1 | 696 |
| BP_S_m | 27 | 2 | 32 | 2 | 2 | 94 | 29 | 85 | 1 | 76 | 3 | 25 | 1 | 17 | 15 | 21 | 5 | 4 | 1 | 152 | 11 | 27 | 2 | 7 | 33 | 6 | 1 | 142 | 1 | 186 |
| BP_S_d | 53 | 1 | 2 | 2 | 6 | 113 | 10 | 2 | 5 | 33 | 4 | 1 | 67 | 6 | 1 | 3 | 1 | 2 | 1 | 80 | 59 | 14 | 2 | 1 | 40 | 16 | 2 | 1 | 20 | 2 |
| Arkona_N_s | 91 | 12 | 43 | 32 | 5 | 65 | 2 | 36 | 6 | 26 | 1 | 6 | 4 | 54 | 67 | 20 | 2 | 11 | 4 | 103 | 6 | 1 | 3 | 1 | 3 | 1 | 2 | 100 | 1 | 729 |
| Arkona_N_m | 29 | 3 | 4 | 1 | 5 | 12 | 3 | 13 | 2 | 18 | 1 | 1 | 2 | 4 | 6 | 1 | 8 | 1 | 2 | 1 | 4 | 2 | 6 | 1 | 10 | 1 | 4 | 52 | 2 | 188 |
| Arkona_N_d | 124 | 4 | 11 | 2 | 4 | 13 | 1 | 17 | 2 | 7 | 1 | 1 | 17 | 24 | 2 | 3 | 3 | 4 | 1 | 6 | 4 | 2 | 2 | 1 | 8 | 1 | 5 | 25 | 4 | 151 |
| Arkona_S_s | 75 | 4 | 119 | 23 | 8 | 3 | 2 | 13 | 3 | 21 | 1 | 3 | 2 | 165 | 60 | 27 | 4 | 5 | 5 | 16 | 5 | 3 | 1 | 1 | 8 | 1 | 1 | 75 | 2 | 54 |
| Arkona_S_m | 55 | 2 | 30 | 3 | 8 | 11 | 2 | 52 | 6 | 8 | 2 | 4 | 4 | 18 | 4 | 5 | 1 | 2 | 2 | 3 | 1 | 37 | 19 | 1 | 3 | 2 | 4 | 20 | 2 | 305 |
| Arkona_S_d | 24 | 2 | 1 | 2 | 1 | 1 | 4 | 9 | 1 | 7 | 4 | 4 | 7 | 5 | 4 | 1 | 4 | 1 | 2 | 13 | 2 | 11 | 1 | 1 | 1 | 2 | 2 | 6 | 2 | 16 |
| Oresund_s | 51 | 1 | 85 | 4 | 16 | 3 | 3 | 82 | 6 | 3 | 2 | 2 | 1 | 89 | 14 | 26 | 3 | 1 | 1 | 272 | 1 | 1 | 3 | 2 | 1 | 2 | 1 | 68 | 2 | 187 |
| Oresund_m | 112 | 2 | 34 | 3 | 18 | 1 | 1 | 48 | 2 | 7 | 3 | 2 | 1 | 37 | 2 | 7 | 2 | 1 | 2 | 397 | 4 | 4 | 1 | 2 | 1 | 2 | 1 | 33 | 6 | 128 |
| Oresund_d | 72 | 3 | 14 | 1 | 3 | 2 | 1 | 6 | 3 | 3 | 2 | 1 | 2 | 4 | 1 | 43 | 1 | 2 | 9 | 13 | 2 | 16 | 118 | 2 | 1 | 194 | 1 | 1 | 2 | 2 |
| Skagerrak_s | 1 | 1 | 21 | 1 | 2 | 1 | 11 | 3 | 1 | 2 | 1 | 1 | 2 | 9 | 1 | 2 | 20 | 2 | 1 | 3 | 1 | 2 | 28 | 1 | 12 | 1 | 1 | 2 | 2 | 272 |
| Skagerrak_m | 215 | 1 | 1 | 2 | 1 | 1 | 1 | 1 | 1 | 2 | 1 | 1 | 58 | 1 | 2 | 3 | 1 | 1 | 2 | 10 | 2 | 4 | 2 | 3 | 1 | 1 | 1 | 1 | 1 | 2 |
| Skagerrak_d | 1 | 1 | 1 | 1 | 2 | 2 | 2 | 1 | 1 | 2 | 1 | 2 | 128 | 1 | 1 | 2 | 1 | 1 | 1 | 1 | 1 | 1 | 68 | 1 | 1 | 1 | 1 | 1 | 3 | 1 |
| 120314 | 2 | 0 | 2 | 0 | 0 | 10 | 0 | 0 | 2 | 7 | 3 | 0 | 9 | 6 | 0 | 4 | 0 | 1 | 0 | 11 | 0 | 0 | 1 | 0 | 9 | 1 | 3 | 0 | 1 | 3 |
| 120328 | 3 | 0 | 2 | 0 | 0 | 9 | 2 | 4 | 0 | 9 | 7 | 0 | 0 | 6 | 0 | 2 | 0 | 0 | 0 | 4 | 1 | 1 | 0 | 0 | 0 | 0 | 1 | 0 | 3 | 5 |
| 120403 | 19 | 4 | 5 | 1 | 1 | 5 | 0 | 0 | 1 | 0 | 17 | 1 | 4 | 7 | 1 | 1 | 1 | 1 | 1 | 3 | 5 | 1 | 2 | 2 | 2 | 2 | 1 | 1 | 2 | 3 |
| 120416 | 61 | 1 | 5 | 1 | 2 | 7 | 0 | 0 | 3 | 0 | 28 | 4 | 10 | 4 | 4 | 6 | 2 | 5 | 3 | 24 | 26 | 11 | 7 | 1 | 6 | 2 | 6 | 9 | 13 | 1 |
| 120516 | 44 | 8 | 16 | 2 | 1 | 4 | 0 | 0 | 3 | 0 | 1 | 4 | 1 | 13 | 1 | 7 | 1 | 9 | 7 | 11 | 1 | 7 | 2 | 3 | 1 | 3 | 1 | 4 | 2 | 4 |
| 120531 | 40 | 4 | 32 | 1 | 1 | 4 | 0 | 0 | 1 | 0 | 33 | 2 | 1 | 15 | 3 | 16 | 1 | 1 | 1 | 54 | 4 | 5 | 2 | 1 | 1 | 1 | 1 | 30 | 2 | 8 |
| 120604 | 6 | 2 | 28 | 2 | 2 | 6 | 0 | 0 | 1 | 0 | 1 | 1 | 1 | 2 | 2 | 9 | 2 | 5 | 4 | 3 | 8 | 1 | 4 | 4 | 2 | 1 | 1 | 8 | 2 | 1 |
| 120613 | 24 | 1 | 9 | 2 | 3 | 5 | 0 | 0 | 2 | 0 | 1 | 2 | 2 | 14 | 4 | 24 | 3 | 1 | 3 | 60 | 8 | 2 | 2 | 1 | 4 | 1 | 1 | 30 | 2 | 21 |
| 120619 | 236 | 41 | 41 | 11 | 12 | 1 | 0 | 0 | 1 | 0 | 1 | 10 | 1 | 56 | 72 | 46 | 1 | 1 | 1 | 237 | 2 | 21 | 1 | 1 | 2 | 2 | 2 | 48 | 1 | 40 |
| 120628 | 64 | 22 | 17 | 2 | 15 | 3 | 0 | 0 | 2 | 0 | 9 | 2 | 2 | 9 | 7 | 7 | 1 | 1 | 3 | 50 | 1 | 17 | 2 | 6 | 4 | 1 | 3 | 6 | 3 | 1 |
| 120705 | 100 | 150 | 6 | 16 | 96 | 2 | 0 | 0 | 33 | 0 | 2 | 2 | 1 | 7 | 4 | 2 | 11 | 7 | 1 | 75 | 5 | 6 | 5 | 6 | 2 | 1 | 7 | 1 | 2 | 3 |
| 120709 | 9 | 9 | 5 | 6 | 4 | 3 | 0 | 0 | 7 | 0 | 2 | 2 | 1 | 7 | 18 | 2 | 1 | 1 | 2 | 3 | 3 | 6 | 2 | 1 | 2 | 1 | 1 | 1 | 1 | 1 |
| 120717 | 4 | 27 | 6 | 4 | 3 | 3 | 0 | 0 | 1 | 0 | 1 | 3 | 2 | 3 | 6 | 4 | 3 | 1 | 2 | 1 | 4 | 2 | 3 | 5 | 1 | 8 | 1 | 4 | 2 | 9 |
| 120802 | 16 | 60 | 7 | 8 | 32 | 1 | 0 | 0 | 8 | 0 | 4 | 22 | 1 | 6 | 4 | 2 | 2 | 7 | 1 | 13 | 1 | 2 | 1 | 1 | 1 | 1 | 1 | 1 | 1 | 4 |
| 120806 | 9 | 30 | 8 | 1 | 4 | 2 | 0 | 0 | 6 | 0 | 2 | 13 | 15 | 7 | 2 | 3 | 8 | 4 | 2 | 2 | 4 | 7 | 4 | 4 | 2 | 1 | 3 | 2 | 2 | 2 |
| 120813 | 0 | 0 | 13 | 0 | 0 | 0 | 0 | 0 | 0 | 0 | 0 | 0 | 0 | 0 | 0 | 0 | 0 | 0 | 0 | 0 | 0 | 0 | 0 | 0 | 0 | 0 | 0 | 0 | 0 | 0 |
| 120820 | 70 | 200 | 13 | 14 | 80 | 1 | 0 | 0 | 32 | 0 | 3 | 1 | 2 | 9 | 20 | 2 | 5 | 9 | 1 | 8 | 2 | 1 | 2 | 1 | 2 | 1 | 6 | 94 | 2 | 11 |
| 120823 | 81 | 288 | 14 | 69 | 128 | 22 | 0 | 0 | 7 | 0 | 44 | 1 | 2 | 5 | 30 | 3 | 26 | 7 | 28 | 23 | 1 | 2 | 10 | 1 | 2 | 1 | 73 | 2 | 1 | 23 |
| 120828 | 62 | 87 | 5 | 19 | 133 | 4 | 0 | 0 | 1 | 0 | 20 | 5 | 1 | 10 | 14 | 28 | 18 | 3 | 18 | 2 | 13 | 1 | 1 | 5 | 1 | 1 | 2 | 2 | 1 | 10 |
| 120903 | 27 | 99 | 6 | 5 | 38 | 3 | 0 | 0 | 7 | 0 | 2 | 3 | 1 | 5 | 6 | 4 | 7 | 4 | 2 | 2 | 1 | 1 | 2 | 1 | 2 | 1 | 1 | 1 | 1 | 10 |
| 120920 | 248 | 166 | 34 | 38 | 109 | 20 | 0 | 0 | 12 | 0 | 98 | 2 | 26 | 36 | 32 | 8 | 6 | 4 | 2 | 66 | 7 | 10 | 2 | 2 | 2 | 6 | 4 | 2 | 8 | 10 |
| 120924 | 119 | 80 | 19 | 4 | 45 | 29 | 0 | 0 | 13 | 0 | 14 | 1 | 1 | 6 | 42 | 5 | 15 | 31 | 24 | 4 | 3 | 1 | 75 | 1 | 1 | 1 | 41 | 2 | 1 | 28 |
| 121001 | 204 | 185 | 14 | 13 | 90 | 15 | 0 | 0 | 13 | 0 | 47 | 1 | 11 | 12 | 10 | 10 | 10 | 1 | 7 | 49 | 4 | 4 | 10 | 1 | 1 | 2 | 24 | 2 | 2 | 10 |
| 121004 | 21 | 37 | 12 | 8 | 15 | 3 | 0 | 0 | 6 | 0 | 8 | 6 | 3 | 5 | 2 | 4 | 4 | 6 | 4 | 19 | 1 | 1 | 3 | 1 | 1 | 5 | 1 | 3 | 1 | 1 |
| 121015 | 153 | 227 | 19 | 18 | 127 | 3 | 0 | 0 | 18 | 0 | 38 | 1 | 4 | 9 | 8 | 3 | 5 | 2 | 14 | 39 | 3 | 10 | 6 | 1 | 2 | 2 | 65 | 1 | 2 | 2 |
| 121022 | 67 | 75 | 13 | 36 | 87 | 1 | 0 | 0 | 4 | 0 | 16 | 1 | 2 | 10 | 6 | 2 | 2 | 2 | 2 | 6 | 3 | 4 | 6 | 1 | 1 | 1 | 12 | 1 | 1 | 11 |
| 121028 | 5 | 35 | 2 | 14 | 6 | 3 | 0 | 0 | 8 | 0 | 1 | 1 | 1 | 5 | 4 | 5 | 7 | 3 | 2 | 4 | 1 | 1 | 2 | 1 | 1 | 2 | 1 | 2 | 1 | 6 |
| 121105 | 53 | 63 | 17 | 8 | 6 | 9 | 0 | 0 | 9 | 0 | 3 | 1 | 8 | 9 | 8 | 13 | 4 | 3 | 10 | 1 | 3 | 2 | 12 | 1 | 6 | 4 | 19 | 2 | 1 | 2 |
| 121128 | 45 | 63 | 11 | 20 | 71 | 3 | 0 | 0 | 10 | 0 | 33 | 1 | 14 | 21 | 2 | 2 | 4 | 2 | 3 | 35 | 2 | 5 | 8 | 1 | 1 | 2 | 24 | 2 | 7 | 4 |
| 121220 | 121 | 20 | 13 | 7 | 39 | 19 | 0 | 0 | 17 | 0 | 64 | 1 | 38 | 82 | 1 | 4 | 1 | 4 | 1 | 185 | 43 | 9 | 7 | 1 | 1 | 13 | 6 | 1 | 26 | 3 |

**Supplementary Table 4**. Breadth per sample and genome. Values equal to or greater than 40% as bold.

| Sample | BACL1 | BACL2 | BACL3 | BACL4 | BACL5 | BACL6 | BACL7 | BACL8 | BACL9 | BACL10 | BACL11 | BACL12 | BACL13 | BACL14 | BACL15 | BACL16 | BACL17 | BACL18 | BACL19 | BACL20 | BACL21 | BACL22 | BACL23 | BACL24 | BACL25 | BACL26 | BACL27 | BACL28 | BACL29 | BACL30 |
|---|---|---|---|---|---|---|---|---|---|---|---|---|---|---|---|---|---|---|---|---|---|---|---|---|---|---|---|---|---|---|
| BB_s | 5 | **90** | 1 | **87** | 1 | **88** | **91** | 2 | **67** | **88** | 3 | 1 | 37 | 12 | **87** | 14 | 11 | **53** | **85** | 16 | **54** | 6 | 2 | 0 | **40** | 0 | **90** | 11 | **66** | **86** |
| BB_m | 34 | **84** | 7 | **89** | 19 | **83** | **90** | 8 | **88** | **89** | **76** | 7 | **89** | 20 | **87** | **60** | 15 | **80** | **89** | **74** | **89** | 13 | 9 | 2 | **54** | 3 | **87** | 31 | **76** | **88** |
| BB_d | **41** | **81** | 3 | **89** | 21 | **76** | **90** | 4 | **89** | **90** | **81** | 8 | **89** | 21 | **85** | **57** | 13 | **81** | **89** | **70** | **89** | 12 | 11 | 3 | **59** | 5 | **88** | 30 | **68** | **85** |
| BS_s | **89** | **90** | **60** | **89** | **81** | **75** | **66** | **58** | **56** | **90** | **62** | 3 | 28 | **81** | **89** | **87** | **91** | **42** | **90** | **83** | **76** | **80** | 3 | 27 | **82** | 25 | **86** | **90** | **74** | **81** |
| BS_m | **89** | **82** | **74** | **90** | **88** | **81** | **90** | **86** | **79** | **90** | **89** | 10 | **89** | **90** | **90** | **88** | **91** | **80** | **90** | **86** | **89** | **83** | 9 | **70** | **80** | **89** | **83** | **90** | **88** | **79** |
| BS_d | **83** | **82** | **58** | **90** | **89** | **74** | **85** | **88** | **90** | **80** | **90** | 11 | **82** | **89** | **89** | **90** | **59** | **84** | **89** | **82** | **90** | **84** | 19 | **83** | **83** | **90** | **90** | **91** | **89** | **90** |
| BP_N_s | **88** | **83** | **90** | **90** | **87** | **76** | **90** | **90** | 39 | **90** | **41** | **90** | 4 | **90** | **77** | **91** | **91** | **90** | **84** | **83** | **90** | **87** | **64** | 0 | **40** | 32 | **90** | **79** | **75** | **81** |
| BP_N_m | **65** | 6 | 8 | 2 | 2 | **73** | **75** | 19 | **44** | **89** | **64** | 6 | 15 | **65** | 3 | 8 | 11 | 2 | 18 | **83** | **88** | **69** | 1 | 16 | **81** | **80** | **56** | 19 | **82** | **87** |
| BP_N_d | **78** | 30 | 8 | 9 | 5 | **81** | **90** | 38 | **81** | **89** | **85** | 2 | **87** | **84** | 9 | 12 | 4 | 2 | 7 | **88** | **90** | 19 | 2 | **45** | **90** | **89** | 27 | 13 | **89** | **86** |
| BP_E_s | **76** | **87** | **90** | **69** | 10 | **89** | **91** | **88** | 16 | **89** | 9 | 19 | 4 | **88** | **90** | **90** | **90** | **87** | **90** | **46** | **90** | **85** | 11 | 0 | **85** | 25 | **90** | **76** | **69** | **77** |
| BP_E_m | 38 | 8 | 10 | 2 | 3 | **90** | **90** | 34 | 19 | **89** | 34 | **58** | 3 | **73** | 1 | 12 | 1 | 1 | 2 | **88** | **87** | **77** | 2 | 7 | **79** | **90** | 6 | 8 | **83** | **91** |
| BP_E_d | **75** | 36 | 11 | 12 | 4 | **82** | **72** | 13 | **85** | **86** | 7 | 3 | **85** | **76** | 9 | 10 | 3 | 1 | 14 | **82** | **76** | 3 | 1 | 4 | **90** | **81** | 38 | 19 | **65** | **65** |
| BP_W_s | **80** | **67** | **88** | 34 | 4 | **87** | **68** | **54** | 14 | **83** | 1 | **80** | 2 | **77** | **90** | **79** | **90** | **89** | **42** | **68** | **59** | 35 | **43** | 0 | **62** | 2 | **73** | **81** | 38 | **80** |
| BP_W_m | **78** | 27 | 25 | 8 | 5 | **80** | **90** | **41** | **84** | **86** | **87** | 11 | **89** | **77** | 7 | 18 | 4 | 4 | 13 | **87** | **90** | **61** | 3 | 1 | **79** | **89** | **44** | 14 | **85** | **87** |
| BP_W_d | **80** | 28 | 28 | 9 | 5 | **79** | **89** | **57** | **86** | **84** | **88** | 16 | **83** | **89** | 7 | 21 | 4 | 5 | 12 | **89** | **90** | **76** | 4 | 1 | **75** | **90** | **46** | 17 | **90** | **89** |
| BP_S_s | **81** | **89** | **78** | **74** | **48** | **88** | **90** | **81** | **49** | **89** | 8 | **90** | 11 | **80** | **88** | **89** | **90** | **90** | 30 | **82** | **91** | **82** | 22 | 1 | **67** | 30 | **63** | **80** | **77** | **79** |
| BP_S_m | **89** | **69** | **90** | 36 | 13 | **75** | **90** | **83** | 31 | **90** | **52** | **90** | 4 | **87** | **90** | **89** | **92** | **81** | 12 | **85** | **91** | **82** | 21 | 0 | **90** | **61** | 33 | **83** | **85** | **81** |
| BP_S_d | **88** | 5 | 31 | 2 | **56** | **73** | **80** | **89** | **82** | **90** | **74** | 2 | **84** | **87** | 3 | **44** | 4 | 1 | 25 | **82** | **90** | 26 | 1 | 1 | **90** | **90** | 22 | **87** | **89** | **87** |
| Arkona_N_s | **82** | **89** | **90** | **87** | **66** | **87** | **66** | **90** | **88** | **90** | 25 | **90** | 12 | **88** | **90** | **88** | **91** | **77** | **52** | **84** | **88** | **73** | 14 | 0 | **91** | 32 | **70** | **75** | **85** | **80** |
| Arkona_N_m | **88** | **54** | **82** | 33 | **56** | **90** | 18 | **89** | 17 | **88** | 4 | 21 | **44** | **85** | **86** | 27 | **90** | 20 | **48** | **87** | **68** | **44** | 12 | 0 | **90** | 13 | **44** | **90** | **87** | **81** |
| Arkona_N_d | **84** | **56** | **89** | 37 | **81** | **90** | 13 | **90** | 19 | **89** | 5 | 13 | **76** | **86** | **84** | **49** | **81** | 28 | 32 | **89** | **77** | 37 | 24 | 1 | **92** | 30 | 30 | **90** | **89** | **79** |
| Arkona_S_s | **84** | **88** | **79** | **86** | **87** | **90** | 9 | **90** | **89** | **89** | 3 | **90** | 8 | **80** | **89** | **90** | **91** | **76** | 38 | **86** | **79** | **50** | **49** | 1 | **83** | 6 | 27 | **84** | **81** | **83** |
| Arkona_S_m | **88** | **43** | **90** | 27 | **81** | **89** | 7 | **90** | 20 | **89** | 3 | **46** | 22 | **86** | **76** | **87** | **60** | 20 | 11 | **88** | **68** | 15 | 27 | 0 | **86** | 15 | 9 | **90** | **86** | **81** |
| Arkona_S_d | **87** | 11 | **84** | 4 | **43** | **73** | 3 | **83** | 4 | **82** | 1 | 2 | **47** | **81** | 12 | 22 | 11 | 3 | 2 | **81** | 27 | 3 | 3 | 0 | **63** | 10 | 6 | **84** | **84** | **90** |
| Oresund_s | **81** | **71** | **78** | **56** | **89** | **59** | 3 | **88** | **56** | **88** | 2 | **73** | 7 | **78** | **90** | **89** | **67** | 39 | 9 | **83** | **65** | 22 | **79** | 1 | **53** | 6 | 14 | **89** | **87** | **81** |
| Oresund_m | **83** | **56** | **84** | **42** | **88** | **50** | 8 | **88** | 38 | **88** | 4 | **58** | 24 | **81** | **90** | **88** | **51** | 27 | 5 | **86** | **69** | 18 | **54** | 1 | **51** | 9 | 6 | **90** | **87** | **80** |
| Oresund_d | **85** | 1 | **88** | 0 | **49** | 15 | 3 | **86** | 2 | 36 | 2 | 2 | **63** | **85** | 4 | 18 | 2 | 2 | 0 | **87** | **58** | 4 | 1 | 3 | 9 | 9 | 0 | **64** | **87** | **83** |
| Skagerrak_s | **79** | 1 | **88** | 0 | 38 | 5 | 1 | **78** | 0 | 8 | 0 | 0 | 5 | **84** | 1 | 5 | 0 | 0 | 0 | **74** | 4 | 3 | 0 | 7 | 6 | 1 | 0 | 35 | **88** | 10 |
| Skagerrak_m | 27 | 3 | 26 | 1 | 9 | 13 | 2 | 5 | 0 | 3 | 2 | 0 | **66** | 13 | 4 | 4 | 0 | 0 | 0 | 30 | 4 | 1 | 0 | 7 | 2 | 3 | 1 | 10 | **89** | 20 |
| Skagerrak_d | 9 | 3 | 7 | 1 | 5 | 38 | 1 | 2 | 0 | 3 | 1 | 0 | **65** | 5 | 1 | 2 | 0 | 0 | 0 | 23 | 1 | 1 | 0 | 2 | 10 | 1 | 2 | 1 | **89** | 9 |
| 120314 | **83** | 27 | **79** | 5 | 17 | **91** | **85** | 31 | **70** | **90** | **83** | 5 | **90** | **81** | 7 | **65** | 30 | **57** | **50** | **87** | **51** | 3 | **48** | 1 | **93** | **54** | **73** | 6 | 28 | **79** |
| 120328 | **81** | 34 | **56** | 12 | 20 | **91** | **86** | **89** | 19 | **90** | **90** | 4 | **87** | **89** | 7 | **62** | 12 | 12 | 14 | **88** | **66** | 15 | 7 | 1 | **60** | **43** | **48** | 7 | **67** | **90** |
| 120403 | **86** | **43** | 39 | 18 | 24 | **88** | **90** | **58** | 19 | **90** | **90** | 2 | **90** | **88** | 6 | 35 | 5 | 15 | 8 | **89** | **91** | 21 | 13 | 0 | **83** | **90** | 26 | 4 | **84** | **40** |
| 120416 | **87** | **55** | **60** | 17 | 28 | **90** | **90** | **83** | 10 | **90** | **89** | 1 | **72** | **87** | 19 | **48** | 17 | 15 | 8 | **89** | **87** | **56** | 9 | 4 | **61** | **87** | 38 | 21 | **88** | **40** |
| 120516 | **87** | **48** | **91** | 25 | 14 | **93** | **90** | **90** | 18 | **89** | **61** | 3 | 7 | **92** | **63** | **90** | 7 | 16 | 2 | **90** | **90** | **70** | 6 | **40** | **89** | **91** | 8 | **90** | **58** | **66** |
| 120531 | **86** | **46** | **83** | 13 | 12 | **67** | **77** | **88** | 6 | **88** | 19 | 3 | 1 | **82** | **43** | **66** | 6 | 12 | 1 | **86** | **60** | 18 | 3 | 1 | **43** | 27 | 4 | **78** | 14 | 19 |
| 120604 | **89** | **68** | **90** | 33 | 25 | **82** | **46** | **84** | 7 | **89** | 3 | **56** | 0 | **90** | **93** | **90** | 24 | 6 | 2 | **90** | **74** | **79** | 2 | 0 | 20 | **45** | 7 | **90** | 9 | **90** |
| 120613 | **66** | 26 | **89** | 6 | 3 | **72** | **65** | **89** | 3 | **83** | 3 | 3 | 0 | **67** | **40** | **89** | 10 | 15 | 1 | **58** | **54** | 9 | 3 | 0 | **41** | 12 | 3 | **86** | 1 | **79** |
| 120619 | **90** | **81** | **91** | **61** | **46** | **91** | **44** | **90** | 9 | **89** | 14 | **69** | 1 | **89** | **91** | **90** | **40** | 32 | 2 | **88** | **47** | **80** | 5 | 0 | 25 | 14 | 14 | **90** | 10 | **90** |
| 120628 | **81** | **90** | **90** | **90** | **90** | **68** | **62** | **90** | 20 | **88** | 19 | **90** | 3 | **90** | **90** | **90** | **53** | 31 | 5 | **82** | **81** | **90** | 7 | 0 | 5 | **66** | 32 | **90** | 5 | **90** |
| 120705 | **89** | **88** | **89** | **69** | **73** | **79** | **51** | **87** | 6 | **85** | **86** | 37 | 0 | **89** | **89** | **90** | **40** | 21 | 2 | **89** | **74** | **79** | **66** | 0 | 4 | 6 | 14 | **69** | 4 | **58** |
| 120709 | **87** | **83** | **76** | **89** | **81** | 39 | 4 | **73** | **89** | 21 | **87** | 1 | 1 | **82** | **88** | 7 | **89** | **74** | **73** | **73** | 11 | 11 | **57** | 0 | 0 | 0 | **86** | 7 | 1 | **72** |
| 120717 | 0 | **89** | **84** | **73** | **47** | 34 | 1 | 38 | 37 | **70** | 14 | **80** | 0 | **74** | **89** | 15 | **63** | 18 | 4 | **74** | 8 | **50** | 26 | 0 | 0 | 0 | 20 | 28 | 0 | **47** |
| 120802 | **69** | **88** | **71** | **67** | **63** | 26 | 1 | 15 | **46** | 37 | 9 | 36 | 0 | **72** | **86** | 5 | 34 | **70** | 4 | **68** | 5 | 16 | 13 | 0 | 0 | 0 | 10 | 7 | 0 | **48** |
| 120806 | **89** | **90** | **91** | **89** | **90** | 29 | 4 | **62** | **90** | **55** | **68** | **90** | 0 | **89** | **91** | 8 | **57** | **85** | 15 | **86** | 13 | **48** | 12 | 1 | 1 | 0 | 28 | 8 | 0 | **90** |
| 120813 | **86** | **89** | **74** | **73** | **86** | 26 | 2 | 26 | **83** | 22 | 23 | **90** | 0 | **83** | **73** | 5 | 33 | **55** | 7 | **73** | 3 | 13 | 4 | 0 | 1 | 0 | 15 | 6 | 0 | **73** |
| 120820 | **89** | **83** | **90** | **90** | **89** | **43** | 8 | **87** | **87** | **61** | **47** | **90** | 0 | **88** | **90** | 15 | **89** | **89** | **53** | **76** | 18 | **47** | **78** | 0 | 3 | 0 | **84** | 19 | 0 | **90** |
| 120823 | **88** | **82** | **90** | **90** | **87** | 39 | 4 | **93** | **90** | **64** | **91** | 25 | 0 | **89** | **90** | 13 | **91** | **90** | **78** | **86** | **66** | **69** | **71** | 0 | 5 | 0 | **88** | 14 | 0 | **91** |
| 120828 | **76** | **82** | **90** | **88** | **84** | **90** | **74** | **90** | **90** | **92** | **90** | 4 | **83** | **89** | **90** | **79** | **90** | **90** | **90** | **90** | **83** | **85** | **90** | 0 | 22 | 14 | **84** | **84** | **83** | **90** |
| 120903 | **86** | **82** | **58** | **86** | **81** | 27 | 4 | **49** | **86** | 16 | **73** | 0 | 1 | **64** | **78** | 5 | **82** | **53** | **53** | **56** | 7 | 6 | 36 | 0 | 1 | 1 | **71** | 5 | 2 | **46** |
| 120920 | **88** | **84** | **57** | **87** | **89** | 27 | 8 | **48** | **87** | 17 | **76** | 0 | 4 | **65** | **79** | 5 | **84** | **53** | **53** | **57** | 8 | 6 | 36 | 0 | 2 | 3 | **72** | 4 | 3 | **45** |
| 120924 | **73** | **72** | **80** | **80** | **75** | **80** | **80** | **80** | **80** | **80** | **71** | 12 | **80** | **80** | **80** | **82** | **82** | **77** | **70** | **75** | **80** | **49** | **82** | 1 | 19 | **79** | **80** | **55** | **79** | **80** |
| 121001 | **88** | **83** | **90** | **90** | **90** | **90** | **92** | **84** | **90** | **89** | **90** | 4 | 26 | **81** | **90** | **88** | **90** | **90** | **90** | **83** | **71** | 31 | **88** | 0 | 20 | 33 | **89** | **84** | 39 | **90** |
| 121004 | **82** | **82** | **90** | **90** | **86** | **89** | **94** | **90** | **90** | **91** | **90** | 4 | **90** | **90** | **90** | **92** | **91** | **92** | **91** | **90** | **85** | **73** | **89** | 0 | 29 | **63** | **90** | **59** | **89** | **90** |
| 121015 | **90** | **90** | **89** | **90** | **79** | **86** | **56** | **78** | **86** | **66** | **88** | 1 | **46** | **84** | **83** | **57** | **71** | **67** | **70** | **73** | 34 | 18 | **87** | 0 | 9 | 19 | **89** | 17 | 30 | **81** |
| 121022 | **82** | **80** | **90** | **90** | **83** | **89** | **79** | **91** | **90** | **91** | **90** | 11 | **85** | **90** | **89** | **76** | **90** | **91** | **90** | **90** | **80** | **88** | **94** | 0 | 10 | 18 | **90** | **47** | **87** | **90** |
| 121028 | **90** | **88** | **90** | **90** | **85** | **75** | **52** | **88** | **94** | **93** | **89** | 9 | **51** | **94** | **92** | **58** | **89** | **79** | **92** | **90** | **59** | **85** | **89** | 0 | 5 | 9 | **90** | 8 | **54** | **93** |
| 121105 | **87** | **90** | **73** | **91** | **88** | **44** | 3 | **77** | **91** | 16 | 25 | 37 | 0 | **60** | **92** | 8 | **91** | **73** | **73** | 39 | 16 | 14 | 36 | 0 | 5 | 0 | **89** | 19 | 1 | **91** |
| 121128 | **85** | **85** | **89** | **87** | **69** | **76** | **78** | **85** | **75** | **80** | **81** | 1 | 34 | **77** | **71** | **62** | **63** | **70** | **71** | **54** | **56** | 35 | **79** | 0 | 4 | 24 | **89** | 9 | 28 | **62** |
| 121220 | **88** | **90** | **92** | **90** | **90** | **91** | **93** | **92** | **90** | **90** | **90** | 3 | **90** | **89** | **79** | **55** | **77** | **73** | **85** | **90** | **89** | **88** | **86** | 1 | 28 | 39 | **90** | 14 | **91** | **90** |
| 121220 | **76** | **90** | **90** | **91** | **90** | **90** | **90** | **90** | **90** | **90** | **87** | 16 | **90** | **85** | **72** | **94** | **90** | **93** | **82** | **83** | **90** | **90** | **94** | 1 | **90** | **90** | **90** | 25 | **90** | **73** |

**Supplementary Table 5**. Correlation (Spearman) of nucleotide diversity for all included BACLs vs. environmental variables. 'Depth' only applicable for samples compared across the vertical dimension. 'Season' only applicable for samples retrieved from the LMO station.

| Baltic Cluster | Sample type | n | Temperature | | | Salinity | | | Chlorophyll a | | | DOC | | | TN | | | Phosphate | | | Nitrite | | | Nitrate | | | Ammonia | | | TIN | | | DON:DIN | | | Depth | | | Oxygen | | | Season (distance) | | |
|---|---|---|---|---|---|---|---|---|---|---|---|---|---|---|---|---|---|---|---|---|---|---|---|---|---|---|---|---|---|---|---|---|---|---|---|---|---|---|---|---|---|---|---|---|
| | | | rho | p | Q | rho | p | Q | rho | p | Q | rho | p | Q | rho | p | Q | rho | p | Q | rho | p | Q | rho | p | Q | rho | p | Q | rho | p | Q | rho | p | Q | rho | p | Q | rho | p | Q | rho | p | Q |
| BACL1 | surf | 5 | 0.67 | 0.219 | 0.892 | 0.97 | 0.005 | 0.158 | -0.05 | 0.935 | 0.972 | 0.36 | 0.553 | 0.954 | 0.05 | 0.935 | 0.972 | -0.70 | 0.188 | 0.876 | 0.71 | 0.182 | 0.871 | 0.29 | 0.638 | 0.921 | 0.30 | 0.624 | 0.961 | 0.40 | 0.505 | 0.949 | -0.40 | 0.505 | 0.969 | | | | -1.00 | 0.001 | 0.001 | | | |
| | depths | 15 | 0.26 | 0.348 | 0.929 | -0.38 | 0.159 | 0.857 | -0.27 | 0.327 | 0.925 | 0.46 | 0.086 | 0.764 | 0.32 | 0.248 | 0.903 | -0.19 | 0.499 | 0.949 | -0.27 | 0.326 | 0.925 | -0.43 | 0.108 | 0.741 | -0.13 | 0.657 | 0.946 | 0.19 | 0.499 | 0.941 | 0.06 | 0.830 | 0.967 | -0.16 | 0.569 | 0.954 | 0.26 | 0.348 | 0.929 | | | |
| | lmo | 20 | -0.82 | 0.001 | 0.001 | 0.35 | 0.125 | 0.825 | -0.62 | 0.004 | 0.131 | -0.63 | 0.003 | 0.101 | | | | | | | | | | 0.41 | 0.076 | 0.923 | | | | | | | | | | | | | | | | 0.08 | 0.188 | 0.768 |
| | surf_lmo | 25 | -0.32 | 0.119 | 0.817 | 0.65 | 0.001 | 0.001 | -0.62 | 0.001 | 0.036 | -0.74 | 0.001 | 0.001 | | | | | | | | | | -0.21 | 0.317 | 0.070 | | | | | | | | | | | | | | | | | | |
| BACL2 | surf | 0 | | | | | | | | | | | | | | | | | | | | | | | | | | | | | | | | | | | | | | | | | | |
| | depths | 0 | | | | | | | | | | | | | | | | | | | | | | | | | | | | | | | | | | | | | | | | | | |
| | lmo | 20 | -0.31 | 0.183 | 0.873 | 0.05 | 0.819 | 0.969 | -0.21 | 0.364 | 0.932 | -0.26 | 0.268 | 0.910 | | | | | | | | | | 0.17 | 0.469 | 0.914 | | | | | | | | | | | | | | | | 0.18 | 0.054 | 0.616 |
| | surf_lmo | 0 | | | | | | | | | | | | | | | | | | | | | | | | | | | | | | | | | | | | | | | | | | |
| BACL3 | surf | 7 | -0.09 | 0.848 | 0.970 | 0.05 | 0.908 | 0.972 | 0.20 | 0.67 | 0.962 | 0.52 | 0.229 | 0.896 | 0.16 | 0.728 | 0.965 | 0.07 | 0.879 | 0.971 | 0.40 | 0.373 | 0.933 | 0.47 | 0.282 | 0.954 | 0.32 | 0.482 | 0.925 | 0.43 | 0.337 | 0.937 | -0.39 | 0.383 | 0.967 | | | | -0.14 | 0.76 | 0.966 | | | |
| | depths | 0 | | | | | | | | | | | | | | | | | | | | | | | | | | | | | | | | | | | | | | | | | | |
| | lmo | 0 | | | | | | | | | | | | | | | | | | | | | | | | | | | | | | | | | | | | | | | | | | |
| | surf_lmo | 0 | | | | | | | | | | | | | | | | | | | | | | | | | | | | | | | | | | | | | | | | | | |
| BACL5 | surf | 0 | | | | | | | | | | | | | | | | | | | | | | | | | | | | | | | | | | | | | | | | | | |
| | depths | 0 | | | | | | | | | | | | | | | | | | | | | | | | | | | | | | | | | | | | | | | | | | |
| | lmo | 13 | 0.07 | 0.831 | 0.970 | 0.22 | 0.475 | 0.947 | 0.21 | 0.481 | 0.948 | 0.04 | 0.887 | 0.971 | | | | | | | | | | 0.18 | 0.552 | 0.530 | | | | | | | | | | | | | | | | -0.18 | 0.940 | 0.986 |
| | surf_lmo | 0 | | | | | | | | | | | | | | | | | | | | | | | | | | | | | | | | | | | | | | | | | | |
| BACL6 | surf | 7 | -0.31 | 0.504 | 0.950 | -0.20 | 0.67 | 0.962 | -0.07 | 0.877 | 0.971 | -0.59 | 0.159 | 0.857 | -0.95 | 0.001 | 0.036 | -0.39 | 0.383 | 0.935 | -0.40 | 0.373 | 0.933 | -0.80 | 0.030 | 0.967 | -0.32 | 0.482 | 0.959 | -0.89 | 0.007 | 0.956 | 0.71 | 0.071 | 0.945 | | | | 0.29 | 0.535 | 0.953 | | | |
| | depths | 20 | -0.13 | 0.594 | 0.957 | 0.06 | 0.80 | 0.968 | -0.17 | 0.488 | 0.948 | -0.33 | 0.170 | 0.865 | -0.08 | 0.74 | 0.965 | -0.02 | 0.943 | 0.973 | 0.22 | 0.370 | 0.933 | -0.40 | 0.091 | 0.873 | 0.12 | 0.622 | 0.876 | -0.14 | 0.576 | 0.970 | 0.18 | 0.459 | 0.950 | 0.16 | 0.505 | 0.956 | 0.03 | 0.909 | 0.972 | | | |
| | lmo | 0 | | | | | | | | | | | | | | | | | | | | | | | | | | | | | | | | | | | | | | | | | | |
| | surf_lmo | 0 | | | | | | | | | | | | | | | | | | | | | | | | | | | | | | | | | | | | | | | | | | |
| BACL8 | surf | 5 | 0.36 | 0.553 | 0.954 | 0.15 | 0.805 | 0.968 | -0.87 | 0.054 | 0.670 | 0.67 | 0.219 | 0.892 | 0.67 | 0.219 | 0.892 | 0.90 | 0.037 | 0.582 | -0.78 | 0.118 | 0.816 | 0.71 | 0.182 | 0.972 | 0.50 | 0.391 | 0.972 | 0.60 | 0.285 | 0.971 | 0.10 | 0.873 | 0.967 | | | | -0.40 | 0.505 | 0.950 | | | |
| | depths | 0 | | | | | | | | | | | | | | | | | | | | | | | | | | | | | | | | | | | | | | | | | | |
| | lmo | 0 | | | | | | | | | | | | | | | | | | | | | | | | | | | | | | | | | | | | | | | | | | |
| | surf_lmo | 0 | | | | | | | | | | | | | | | | | | | | | | | | | | | | | | | | | | | | | | | | | | |
| BACL10 | surf | 0 | | | | | | | | | | | | | | | | | | | | | | | | | | | | | | | | | | | | | | | | | | |
| | depths | 11 | 0.14 | 0.689 | 0.963 | -0.07 | 0.832 | 0.969 | -0.11 | 0.739 | 0.965 | -0.20 | 0.555 | 0.954 | 0.17 | 0.62 | 0.959 | 0.07 | 0.842 | 0.969 | 0.05 | 0.883 | 0.971 | -0.20 | 0.564 | 0.935 | 0.04 | 0.915 | 0.966 | 0.04 | 0.905 | 0.970 | -0.09 | 0.790 | 0.970 | -0.18 | 0.592 | 0.959 | 0.15 | 0.67 | 0.962 | | | |
| | lmo | 0 | | | | | | | | | | | | | | | | | | | | | | | | | | | | | | | | | | | | | | | | | | |
| | surf_lmo | 0 | | | | | | | | | | | | | | | | | | | | | | | | | | | | | | | | | | | | | | | | | | |
| BACL11 | surf | 0 | | | | | | | | | | | | | | | | | | | | | | | | | | | | | | | | | | | | | | | | | | |
| | depths | 0 | | | | | | | | | | | | | | | | | | | | | | | | | | | | | | | | | | | | | | | | | | |
| | lmo | 8 | 0.36 | 0.385 | 0.935 | -0.18 | 0.678 | 0.962 | -0.02 | 0.955 | 0.973 | 0.21 | 0.61 | 0.958 | | | | | | | | | | -0.36 | 0.385 | 0.866 | | | | | | | | | | | | | | | | 0.04 | 0.349 | 0.973 |
| | surf_lmo | 0 | | | | | | | | | | | | | | | | | | | | | | | | | | | | | | | | | | | | | | | | | | |
| BACL14 | surf | 5 | 0.82 | 0.089 | 0.770 | 0.82 | 0.089 | 0.770 | -0.21 | 0.741 | 0.965 | 0.21 | 0.741 | 0.965 | 0.21 | 0.741 | 0.965 | -0.30 | 0.624 | 0.959 | 0.71 | 0.182 | 0.873 | -0.29 | 0.638 | 0.974 | 0.30 | 0.624 | 0.970 | 0.10 | 0.873 | 0.965 | -0.10 | 0.873 | 0.952 | | | | -0.70 | 0.188 | 0.876 | | | |
| | depths | 0 | | | | | | | | | | | | | | | | | | | | | | | | | | | | | | | | | | | | | | | | | | |
| | lmo | 0 | | | | | | | | | | | | | | | | | | | | | | | | | | | | | | | | | | | | | | | | | | |
| | surf_lmo | 0 | | | | | | | | | | | | | | | | | | | | | | | | | | | | | | | | | | | | | | | | | | |
| BACL15 | surf | 5 | 0.56 | 0.322 | 0.924 | -0.15 | 0.805 | 0.968 | -0.67 | 0.219 | 0.892 | -0.56 | 0.322 | 0.924 | 0.56 | 0.322 | 0.924 | 0.80 | 0.104 | 0.796 | | | 0.871 | | | | -0.20 | 0.747 | 0.965 | -0.10 | 0.873 | 0.936 | 0.10 | 0.873 | 0.936 | | | | 0.30 | 0.624 | 0.959 | | | |
| | depths | 0 | | | | | | | | | | | | | | | | | | | | | | | | | | | | | | | | | | | | | | | | | | |
| | lmo | 5 | 0.20 | 0.747 | 0.966 | 0.22 | 0.718 | 0.964 | 0.10 | 0.873 | 0.970 | -0.4 | 0.505 | 0.950 | | | | | | | | | | 0.10 | 0.873 | 0.938 | | | | | | | | | | | | | | | | -0.20 | 0.691 | 0.973 |
| | surf_lmo | 10 | 0.02 | 0.947 | 0.973 | -0.63 | 0.052 | 0.662 | 0.09 | 0.815 | 0.968 | 0.10 | 0.789 | 0.967 | | | | | | | | | | 0.70 | 0.026 | 0.693 | | | | | | | | | | | | | | | | | | |
| BACL20 | surf | 5 | -0.36 | 0.553 | 0.954 | -0.36 | 0.553 | 0.954 | 0.05 | 0.935 | 0.972 | -0.36 | 0.553 | 0.954 | -0.15 | 0.805 | 0.968 | 0.50 | 0.391 | 0.936 | -0.71 | 0.182 | 0.973 | 0.35 | 0.559 | 0.959 | -0.10 | 0.873 | 0.953 | 0.00 | 1.00 | 0.954 | 0.00 | 1.00 | 0.964 | | | | 0.30 | 0.624 | 0.959 | | | |
| | depths | 0 | | | | | | | | | | | | | | | | | | | | | | | | | | | | | | | | | | | | | | | | | | |
| | lmo | 13 | 0.15 | 0.629 | 0.959 | -0.5 | 0.083 | 0.757 | -0.21 | 0.482 | 0.948 | -0.04 | 0.901 | 0.971 | | | | | | | | | | 0.25 | 0.405 | 0.966 | | | | | | | | | | | | | | | | 0.06 | 0.329 | 0.973 |
| | surf_lmo | 18 | -0.13 | 0.607 | 0.958 | -0.4 | 0.104 | 0.796 | -0.10 | 0.705 | 0.964 | 0.26 | 0.299 | 0.918 | | | | | | | | | | 0.45 | 0.060 | 0.974 | | | | | | | | | | | | | | | | | | |
| BACL27 | surf | 0 | | | | | | | | | | | | | | | | | | | | | | | | | | | | | | | | | | | | | | | | | | |
| | depths | 0 | | | | | | | | | | | | | | | | | | | | | | | | | | | | | | | | | | | | | | | | | | |
| | lmo | 5 | 0.90 | 0.037 | 0.582 | -0.58 | 0.308 | 0.921 | 0.50 | 0.391 | 0.936 | 0.70 | 0.188 | 0.876 | | | | | | | | | | -0.30 | 0.624 | 0.971 | | | | | | | | | | | | | | | | 0.88 | 0.008 | 0.182 |
| | surf_lmo | 0 | | | | | | | | | | | | | | | | | | | | | | | | | | | | | | | | | | | | | | | | | | |
| BACL28 | surf | 8 | -0.08 | 0.844 | 0.969 | -0.13 | 0.756 | 0.966 | 0.35 | 0.396 | 0.937 | -0.25 | 0.548 | 0.954 | 0.01 | 0.978 | 0.974 | 0.19 | 0.651 | 0.961 | 0.03 | 0.942 | 0.770 | 0.13 | 0.766 | 0.966 | 0.07 | 0.867 | 0.935 | 0.14 | 0.736 | 0.930 | -0.26 | 0.531 | 0.958 | | | | 0.12 | 0.779 | 0.967 | | | |
| | depths | 13 | 0.04 | 0.894 | 0.971 | -0.06 | 0.837 | 0.969 | 0.08 | 0.788 | 0.967 | 0.00 | 0.993 | 0.974 | 0.10 | 0.740 | 0.965 | -0.05 | 0.859 | 0.970 | -0.09 | 0.778 | 0.973 | 0.04 | 0.896 | 0.964 | 0.19 | 0.534 | 0.960 | 0.18 | 0.553 | 0.969 | 0.11 | 0.721 | 0.950 | -0.18 | 0.546 | 0.956 | 0.07 | 0.831 | 0.969 | | | |
| | lmo | 0 | | | | | | | | | | | | | | | | | | | | | | | | | | | | | | | | | | | | | | | | | | |
| | surf_lmo | 0 | | | | | | | | | | | | | | | | | | | | | | | | | | | | | | | | | | | | | | | | | | |
| BACL30 | surf | 8 | -0.35 | 0.399 | 0.938 | -0.16 | 0.713 | 0.964 | 0.72 | 0.043 | 0.618 | -0.08 | 0.844 | 0.969 | -0.59 | 0.126 | 0.826 | -0.29 | 0.493 | 0.949 | -0.02 | 0.971 | 0.949 | -0.13 | 0.766 | 0.959 | -0.36 | 0.385 | 0.950 | -0.38 | 0.352 | 0.950 | 0.21 | 0.610 | 0.955 | | | | 0.00 | 1.00 | 0.974 | | | |
| | depths | 13 | 0.01 | 0.964 | 0.973 | 0.17 | 0.571 | 0.956 | 0.52 | 0.07 | 0.725 | 0.09 | 0.768 | 0.967 | -0.11 | 0.727 | 0.965 | -0.17 | 0.578 | 0.956 | -0.06 | 0.836 | 0.960 | 0.11 | 0.718 | 0.970 | -0.14 | 0.641 | 0.936 | -0.07 | 0.831 | 0.970 | 0.20 | 0.505 | 0.957 | -0.17 | 0.574 | 0.966 | -0.21 | 0.482 | 0.948 | | | |
| | lmo | 0 | | | | | | | | | | | | | | | | | | | | | | | | | | | | | | | | | | | | | | | | | | |
| | surf_lmo | 0 | | | | | | | | | | | | | | | | | | | | | | | | | | | | | | | | | | | | | | | | | | |

**Supplementary Table 6**. Environmental association analysis showing drivers of population structure per data set and BACL. Spearman correlation of single best predictor and $F_{ST}$ retrieved from Bioenv-package allowing for 1 independent variable (p-values from Mantel's test). Prior to the global RDA analysis up to three independent variables were allowed. For detailed description of analyses, see *Methods, Environmental association analysis.*

| Samples | n | MAG | Single best predictor | rho from bioenv | p-value | *Q*-value | Predictor(s) (3 allowed) | rho from bioenv | Rsq. adj. from RDA | p-value | Conditioned RDA | | |
|---|---|---|---|---|---|---|---|---|---|---|---|---|---|
| Transect surface | 5 | BACL1 | **Salinity** | 0.64 | 0.017 | 0.029 | Salinity+Phosphate+MEM1 | 0.87 | 0.72 | 0.325 | NA | NA | NA |
| Transect surface | 7 | BACL3 | Salinity | 0.64 | 0.059 | 0.075 | **Salinity+DON+MEM2** | 0.89 | 0.98 | 0.001 | **Salinity, r2=0.54, p=0.002** | DON, r2=0.04, p=0.052 | **MEM2, r2=0.06, p=0.037** |
| Transect surface | 7 | BACL6 | **Salinity** | 0.80 | 0.011 | 0.022 | **Temperature+TIN+MEM2** | 0.90 | 0.87 | 0.022 | Temperature, r2=0.02, p=0.223 | **TIN, r2=-0.13, p=0.034** | MEM2, r2=0.07, p=0.146 |
| Transect surface | 5 | BACL8 | Salinity | 0.58 | 0.042 | 0.056 | Salinity+Chla+MEM1 | 0.67 | 0.38 | 0.483 | NA | NA | NA |
| Transect surface | 5 | BACL14 | **Temperature** | 0.76 | 0.008 | 0.019 | **Temperature+TN** | 0.93 | 0.76 | 0.025 | Temperature, r2=0.23, p=0.118 | TN, r2=-0.38, p=0.158 | NA |
| Transect surface | 5 | BACL15 | **Salinity** | 0.83 | 0.025 | 0.038 | not improved | NA | NA | NA | NA | NA | NA |
| Transect surface | 5 | BACL20 | **Salinity** | 0.90 | 0.033 | 0.047 | Temperature+Salinity+Phosphate | 0.91 | 0.99 | 0.008 | Temperature, r2=0.015, p=0.30 | Salinity, r2=0.022, p=0.26 | Phosphate, r2=0.019, p=0.28 |
| Transect surface | 8 | BACL28 | **Temperature** | 0.61 | 0.004 | 0.011 | **Temperature+Salinity+MEM2** | 0.75 | 0.79 | 0.009 | Temperature, r2=-0.01, p=0.413 | Salinity, r2=0.04, p=0.229 | **MEM2, r2=0.27, p=0.041** |
| Transect surface | 8 | BACL30 | **Temperature** | 0.64 | 0.002 | 0.006 | **Salinity+DON:DIN+MEM2** | 0.72 | 0.80 | 0.006 | **Salinity, r2=0.55, p=0.008** | DON:DIN, r2=-0.01, p=0.449 | **MEM2, r2=0.60, p=0.030** |
| LMO | 20 | BACL1 | **Season** | 0.42 | 0.001 | 0.003 | **Temperature+Season** | 0.51 | 0.61 | 0.001 | **Temperature, r2=0.40, p=0.001** | **Season, r2=0.20, p=0.002** | NA |
| LMO | 20 | BACL2 | DOC | 0.25 | 0.109 | 0.125 | DOC+Season | 0.28 | 0.08 | 0.392 | NA | NA | NA |
| LMO | 13 | BACL5 | NO3 | 0.24 | 0.155 | 0.162 | Chl a+DOC+NO3 | 0.33 | 0.04 | 0.351 | NA | NA | NA |
| LMO | 8 | BACL11 | Season | 0.39 | 0.177 | 0.177 | Temperature+Season | 0.41 | 0.47 | 0.129 | NA | NA | NA |
| LMO | 5 | BACL15 | Chl a | 0.58 | 0.100 | 0.120 | **Chl a+NO3** | 0.62 | 0.72 | 0.033 | **Chl a, r2=0.47, p=0.033** | NO3, r2=0.30, p=0.175 | NA |
| LMO | 13 | BACL20 | Temperature | 0.18 | 0.137 | 0.149 | not improved | NA | NA | NA | NA | NA | NA |
| LMO | 5 | BACL27 | **NO3** | 0.82 | 0.025 | 0.038 | not improved | NA | NA | NA | NA | NA | NA |
| Transect surface+LMO | 25 | BACL1 | **Salinity** | 0.59 | 0.001 | 0.003 | **Temperature+MEM1+Season** | 0.60 | 0.62 | 0.001 | **Temperature, r2=0.23, p=0.001** | **MEM1, r2=0.26, p=0.001** | **Season, r2=0.18, p=0.001** |
| Transect surface+LMO | 10 | BACL15 | **Salinity** | 0.62 | 0.001 | 0.003 | **Salinity+MEM1+MEM2** | 0.72 | 0.65 | 0.001 | **Salinity, r2=0.22, p=0.016** | **MEM1, r2=0.43, p=0.006** | MEM2, r2=-0.01, p=0.430 |
| Transect surface+LMO | 18 | BACL20 | **Salinity** | 0.53 | 0.009 | 0.020 | not improved | NA | NA | NA | NA | NA | NA |
| Depths | 15 | BACL1 | **Oxygen** | 0.76 | 0.001 | 0.003 | **Salinity+Oxygen+MEM1** | 0.77 | 0.75 | 0.001 | **Salinity, r2=0.13, p=0.002** | **Oxygen, r2=0.08, p=0.031** | **MEM1, r2=0.08, p=0.015** |
| Depths | 19 | BACL6 | **Salinity** | 0.57 | 0.001 | 0.003 | not improved | NA | NA | NA | NA | NA | NA |
| Depths | 11 | BACL10 | **MEM1** | 0.74 | 0.001 | 0.003 | **Temperature+MEM1** | 0.78 | 0.75 | 0.001 | **Temperature, r2=0.30, p=0.001** | **MEM1, r2=0.44, p=0.004** | NA |
| Depths | 13 | BACL28 | **Oxygen** | 0.51 | 0.001 | 0.003 | not improved | NA | NA | NA | NA | NA | NA |
| Depths | 13 | BACL30 | **Oxygen** | 0.37 | 0.014 | 0.026 | not improved | NA | NA | NA | NA | NA | NA |
